# Guanine-containing ssDNA and RNA induce dimeric and tetrameric SAMHD1 in cryo-EM and binding studies

**DOI:** 10.1101/2023.06.15.544806

**Authors:** Benjamin Orris, Min Woo Sung, Shridhar Bhat, Yingrong Xu, Kevin W. Huynh, Seungil Han, Darren C. Johnson, Benedikt Bosbach, David J. Shields, James T. Stivers

## Abstract

The dNTPase activity of tetrameric SAM and HD domain containing deoxynucleoside triphosphate triphosphohydrolase 1 (SAMHD1) plays a critical role in cellular dNTP regulation. SAMHD1 also associates with stalled DNA replication forks, DNA repair foci, ssRNA, and telomeres. The above functions require nucleic acid binding by SAMHD1, which may be modulated by its oligomeric state. Here we establish that the guanine-specific A1 activator site of each SAMHD1 monomer is used to target the enzyme to guanine nucleotides within single-stranded (ss) DNA and RNA. Remarkably, nucleic acid strands containing a single guanine base induce dimeric SAMHD1, while two or more guanines with ∼20 nucleotide spacing induce a tetrameric form. A cryo-EM structure of ssRNA-bound tetrameric SAMHD1 shows how ssRNA strands bridge two SAMHD1 dimers and stabilize the structure. This ssRNA-bound tetramer is inactive with respect to dNTPase and RNase activity.

## INTRODUCTION

The human enzyme SAMHD1 is a GTP-activated dNTP triphosphohydrolase capable of hydrolyzing all four canonical dNTPs to the deoxynucleoside and triphosphate and plays an important role in nucleotide homeostasis^1, 2^. The complicated activation mechanism of SAMHD1 involves binding of GTP to an activator site (A1) located on each monomer, followed by binding of either dATP, dTTP or dGTP to the second activator site (A2) to drive formation of the dNTPase-active tetramer^3, 4^.

In addition, SAMHD1 has non-dNTPase activities that involve binding of single-stranded (ss) DNA or RNA to the dimer-dimer interface of the tetramer^5, 6^. Interaction of nucleic acids (NA) with the interface is believed to inhibit dNTPase activity by preventing formation of the dNTPase-active tetramer^5^. Recent studies have also shown how phosphorylation of Thr592 (pSAMHD1) may be linked to NA binding through destabilization of the tetramer, allowing rapid access of ssDNA to the dimer interface as compared to the very stable unphosphorylated form of the enzyme^7^. Since pSAMHD1 is implicated in promoting restart of stalled replication forks, one biological function of phosphorylation may be to promote rapid binding to single-stranded nucleic acids (ssNA) present at stalled forks^8^. Additional roles for SAMHD1 that involve ssNA binding have been reported in the context of double-strand break repair^9–12^, class switch recombination^12^, R-loop formation at transcription–replication conflict regions^13^, telomere maintenance^14^, and RNA homeostasis^15^. In addition to ssNA binding, some work has suggested that SAMHD1 has 3′ to 5′ exonuclease activity against ssDNA and ssRNA^16–18^. However, this nuclease activity has been disputed in work from several other labs^5, 6, 19–21^. Nevertheless, a recent study proposed a mechanism where SAMHD1 RNase activity was involved in preventing an overabundance of immunostimulatory cellular ssRNA^15^. A better understanding of nucleic acid binding to SAMHD1 is long overdue given the increasing NA-related activities attributed to the enzyme.

Obtaining high-resolution structural information on SAMHD1-NA complexes has been challenging, but several complementary approaches have provided a low-resolution model^5, 7, 22^. A recent crystal structure of a SAMHD1 dimer in complex with a short 5mer ssDNA strand 5′-C**G**CCT-3′ with non-bridging phosphorothioate internucleotide linkages showed that the A1 site bound to the 5′ guanine nucleotide and suggested that the 3′ end of longer ssDNAs would extend out from the A1 site^22^. This extended binding mode is consistent with a previous ssDNA photochemical cross-linking study that identified a cluster of positively charged residues located on the dimer-dimer interface near the A1 site^5^. As recently suggested^7^, a likely model for binding longer nucleic acids involves two modes: (i) specific binding guanine nucleotides in the A1 site, and (2) non-specific electrostatic interactions with nucleic acid chains using the electropositive dimer surface.

In this study we report the first cryo-EM structures of SAMHD1 bound to ssRNA and perform a comprehensive biochemical characterization of RNA and DNA binding to the enzyme. Surprisingly, the structures reveal that ssNA binding induces both dimeric and tetrameric forms of SAMHD1. A central finding in this work is that exposed guanine bases in ssNA can interact with the A1 site of SAMHD1 and allosterically modulate its oligomeric state. The structures do not support the putative 3′ to 5′ RNA exonuclease activity because the active site is devoid of nucleotides, occluded, and the RNA binding mode does not position the RNA ends towards the catalytic metal center.

## RESULTS

### Contribution of the A1 site to DNA and RNA binding specificity

We designed a large set of oligonucleotides to test various aspects of SAMHD1 binding to single stranded nucleic acids (**Supplemental Table S1**). These constructs probe (i) the binding energy and specificity contributions of the A1 site, (ii) the positional dependence of binding to single dG or G residues in DNA or RNA strands, (iii) the binding contribution of non-specific electrostatic interactions between ssNA and SAMHD1, and (iv) the changes in the oligomeric state of SAMHD1 induced by nucleic acid binding.

To probe the contribution of the A1 site to DNA binding affinity and specificity, we used a mutational approach to switch the specificity of the site from a guanine base to xanthine (X) (**Fig. 1A, B**). This approach is based on a crystal structure (6TX0) showing that the D137N A1 site mutant of SAMHD1 can specifically bind xanthine triphosphate (XTP) in the A1 site^1^. We first established using steady-state kinetic measurements that 0.5 mM XTP induced at least 7-fold greater activation of D137N compared to 0.5 mM GTP and that XTP was a weak activator for wild-type SAMHD1 (**Supplemental Fig. S1**). We then constructed a set of 40mer ssDNA oligonucleotides substituted with single dG, dX or dA residues located two nucleotides from the 5′ end utilizing a 5′ FAM label for anisotropy binding measurements (i.e., 5′FAM-dT**dN**dT_38_) (**Supplemental Table S2, Figure 1**). As observed for binding the nucleotide triphosphates, wild-type SAMHD1 showed a binding selectivity dG > dX > dA ∼ dT (**Fig. 1C**), whereas D137N showed preferential binding in the order dX > dG > dA ∼ dT (**Fig. 1D**). Both control competition experiments using unlabeled 5′dT_40_ and binding assays with the FAM label in the 6-position of dT rather than the terminal 5′ phosphate established that the 5′FAM label has no significant positive or negative impact on ssDNA binding affinity (**Supplemental Figure S2**).

**Figure 1.**
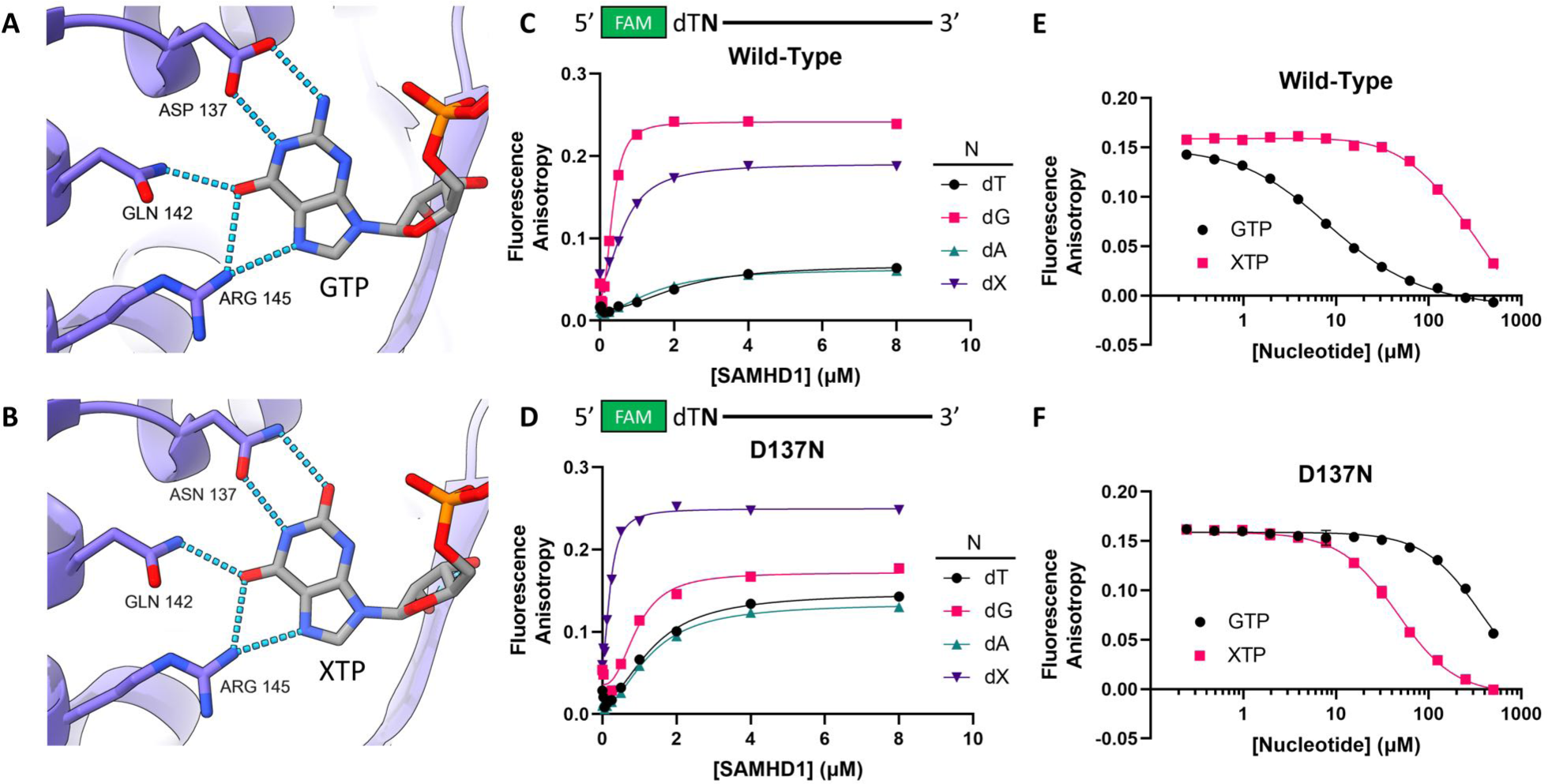
A1 site confers binding specificity for guanine and xanthine bases in ssDNA. (**A**) Guanine-specific hydrogen bonding network formed in the A1 site of wild-type SAMHD1 (PDB: 6TXC). (**B**) Xanthine-specific hydrogen bonding network formed in the A1 site of SAMHD1 D137N (PDB: 6TXA). (**C**) Binding of wild-type SAMHD1 to a series of 5’FAM- labeled 40mer ssDNA oligonucleotides (50 nM) consisting of a dT38 homopolymer with nucleotides (**N**) dT, dG, dA, or dX (deoxyxanthosine) in position two. Error bars indicate standard error of three independent replicate measurements at each SAMHD1 concentration. (**D**) Binding of SAMHD1 D137N to the same series of oligonucleotides as in panel C. Error bars indicate standard error of three independent replicate measurements at each SAMHD1 concentration. Error bars indicate standard error of three independent replicate measurements at each nucleotide concentration. (**E**) Displacement of the dG-containing 40mer (0.5 μM) from wild-type SAMHD1 (1 μM) with GTP (black) or XTP (xanthosine triphosphate, pink). (**F**) Displacement of the dX-containing 40mer (0.5 μM) from SAMHD1 D137N (1 μM) with GTP (black) or XTP (pink). Error bars indicate standard error of three independent replicate measurements at each nucleotide concentration.

As a further measure of the A1 site binding selectivity for the 5′ dG and dX substitutions, we performed competition experiments where XTP and GTP were used to displace 5′FAM-dT**dG**dT_38_ or 5′FAM-dT**dX**dT_38_ from the wild-type and mutant enzymes (**Fig. 1E, F**). Consistent with the anticipated 5′ dG and dX specificities, GTP was a 30-fold better competitor than XTP for wild-type SAMHD1, while XTP was an 8- fold better competitor than GTP for D137N SAMHD1. From these data we conclude that the A1 site in wild-type SAMHD1 encodes binding specificity for G bases in the context of nucleotides and ssDNA. In addition, the specific binding of dX or dG exhibited a 5′ polarity preference, because placement of these bases two nucleotides from the 3′ DNA end (dT_38_**dN**dT-FAM3′) resulted in ∼5-fold weaker binding, highly reduced anisotropy increases, and little dG or dX base specificity as compared to the 5′ constructs (**Supplemental Fig. S3**). The basis for the 5′ G base binding preference is explored further below.

We also performed an analogous binding affinity and specificity study for SAMHD1 in the context of ssRNA using 5′FAM-U**N**U_38_ constructs (**Supplemental Table S2**, where N = G, A, or U) and observed the same 5′ preference for G bases as observed with 5′FAM-dT**dG**dT_38_ (**Supplemental Fig. 4A**).

### Positional effects of single G bases in ssNA

Given the preference of SAMHD1 for binding 5′FAM-dT**dG**dT_38_, we asked how binding would be affected by the position of the dG residue relative to the 5′ end of the ssDNA. We thus constructed nine dT 40mer variants where single dT residues were substituted with dG at positions 1 through 5, 7, 10, 13, 26, and 39 from the 5′ end (**Supplemental Table S2**). DNA anisotropy binding measurements were performed as described above with fitting to the Hill equation (eq 2) (**Fig. 2A**). The salient features of this data set were (i) the *K*_0.5_ values were invariant for dG substitutions in positions 1-7 (average value 0.34 ± 0.02 μM (**Fig. 2B**), (ii) *K*_0.5_ begins to increase with the G residue at the 10^th^ position from the 5′ end (*K*_0.5_ = 0.52 ± 0.05 μM), and (iii) the *K*_0.5_ values plateau after position 26 with an average *K*_0.5_ value of 1.07 ± 0.11 μM, (**Fig. 2B**) which is nearly the same value observed for unsubstituted dT_40_ (*K*_0.5_ = 2.1 ± 0.38 μM). A similar trend was observed with ssRNA 5′FAM-U_40_ variants, where single U residues were substituted with G at positions 13 and 26 from the 5′ end (**Supplemental Table S2, Supplemental Fig. S4B**). Also paralleling the results with DNA, poor binding to an ssRNA strand with a 3′ G was observed (U_38_**G**U-3’FAM) (**Supplemental Table S2**, **Supplemental Fig. 4C**). We conclude that SAMHD1 binds ssDNA and ssRNA using the same A1-site G base recognition mechanism, and that the position of the G base relative to the 5′ end of the NA plays an important role in binding.

**Figure 2.**
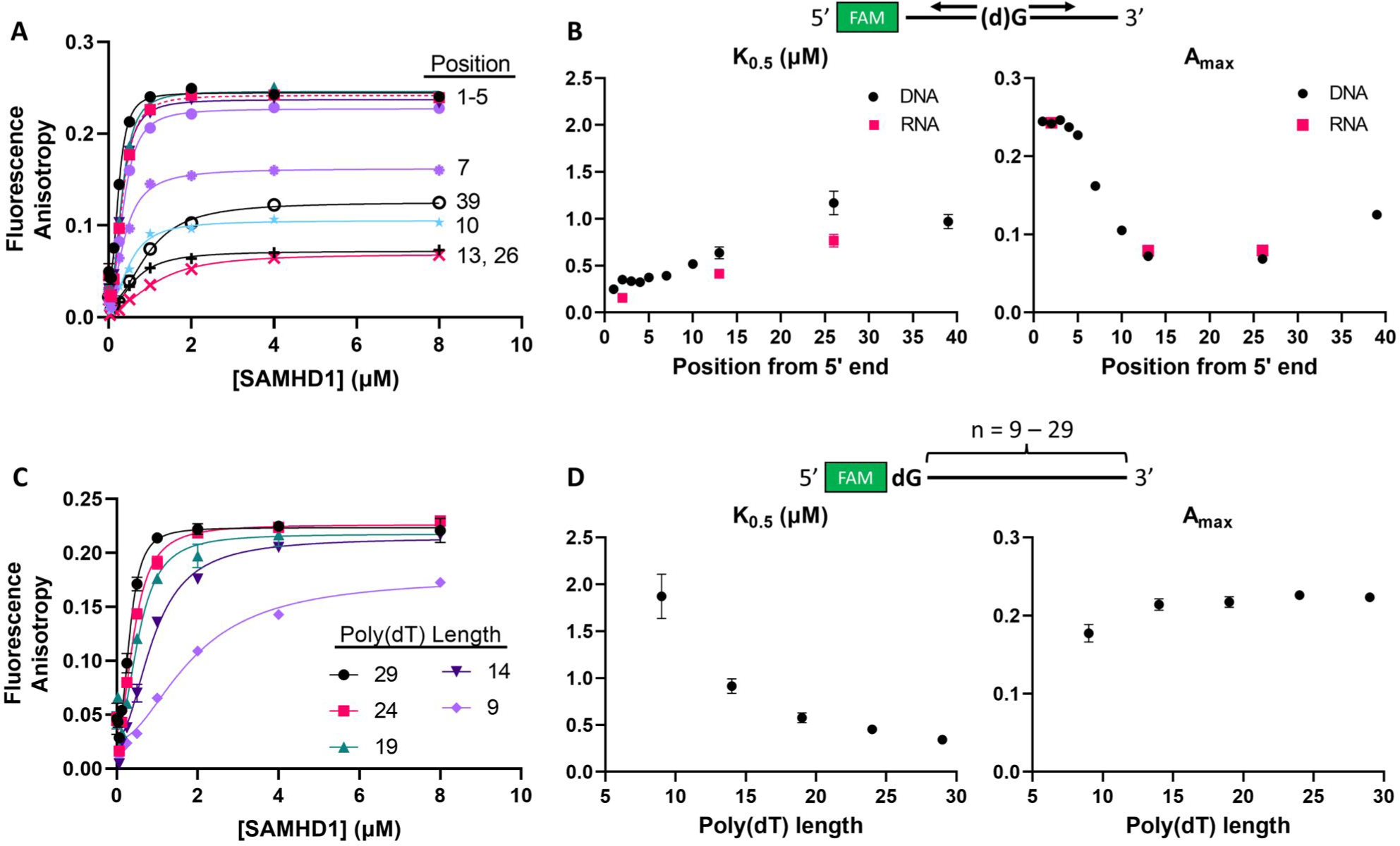
dG positional and oligonucleotide length effects on binding. (**A**) Binding isotherms of SAMHD1 to a series of 5′FAM-labeled ssDNA 40mers containing a single dG nucleotide in varying positions in a dT homopolymer backbone (50 nM each). (**B**) Plot of *K*0.5 (left) and maximum anisotropy (Amax, right) for the ssDNA binding isotherms in (A) (black) and the ssRNA binding isotherms in Supplemental Figure S4B (pink). (**C**) Binding of SAMHD1 to a series of 5′FAM-labeled ssDNA oligonucleotides consisting of a single dG base at the 5′ end followed by dT homopolymer of varying lengths (n = 9 to 29). (**D**) *K*0.5 (left) and maximum anisotropy (Amax, right) observed for the different homopolymer lengths used in panel (C). All Error bars for both *K*0.5 and Amax indicate standard errors as determined by least-squares regression fit to the Hill equation (eq 2).

We then asked whether the binding affinity was impacted by the number of dT nucleotides located 3′ to a single 5′ dG residue (**Supplemental Table S2, Fig. 2C**). We explored binding of five ssDNA constructs where the number of 3′ dT nucleotides was varied in the range 9 to 29 (i.e. 5′FAM-**dG**T_9-29_). Although the binding affinity showed only a modest weakening as the number of 3′ dT nucleotides was decreased from 29 to 19, the binding affinity dropped markedly for the constructs with only nine and fourteen 3′- nucleotides (**Fig. 2D**). Compared to the highest affinity construct (5′FAM-dT**dG**T_29_), 5′FAM-dT**dG**T_9_ bound 5-fold weaker (*K*_0.5_ = 1.87 ± 0.24 μM). We conclude that ssNA binding has two separable contributions (i) preferential A1 site binding of dG residues located within 5 nucleotides from the 5′ NA end, and (ii) non-specific binding of extended nucleic acid strands 3′ to the dG residue, which is optimal when the strand length is at least 25 nucleotides.

### Binding of single G residues in ssDNA or ssRNA induces dimerization of SAMHD1

Since canonical activation of SAMHD1 involves dimerization induced by GTP binding in the A1 site^3, 5, 6^, we reasoned that A1 site mediated binding to guanine-containing ssDNA and ssRNA might also induce dimerization. To address this question, we used our previously validated glutaraldehyde protein crosslinking (GAXL) assay to detect changes in the oligomeric state of SAMHD1 upon binding ssDNA and ssRNA (**Fig. 3**). In this assay, only the protein is crosslinked with glutaraldehyde and less than 5% ssNA remains bound during SDS-PAGE analysis of the crosslinked samples (**Supplemental Fig. S5**).

**Figure 3.**
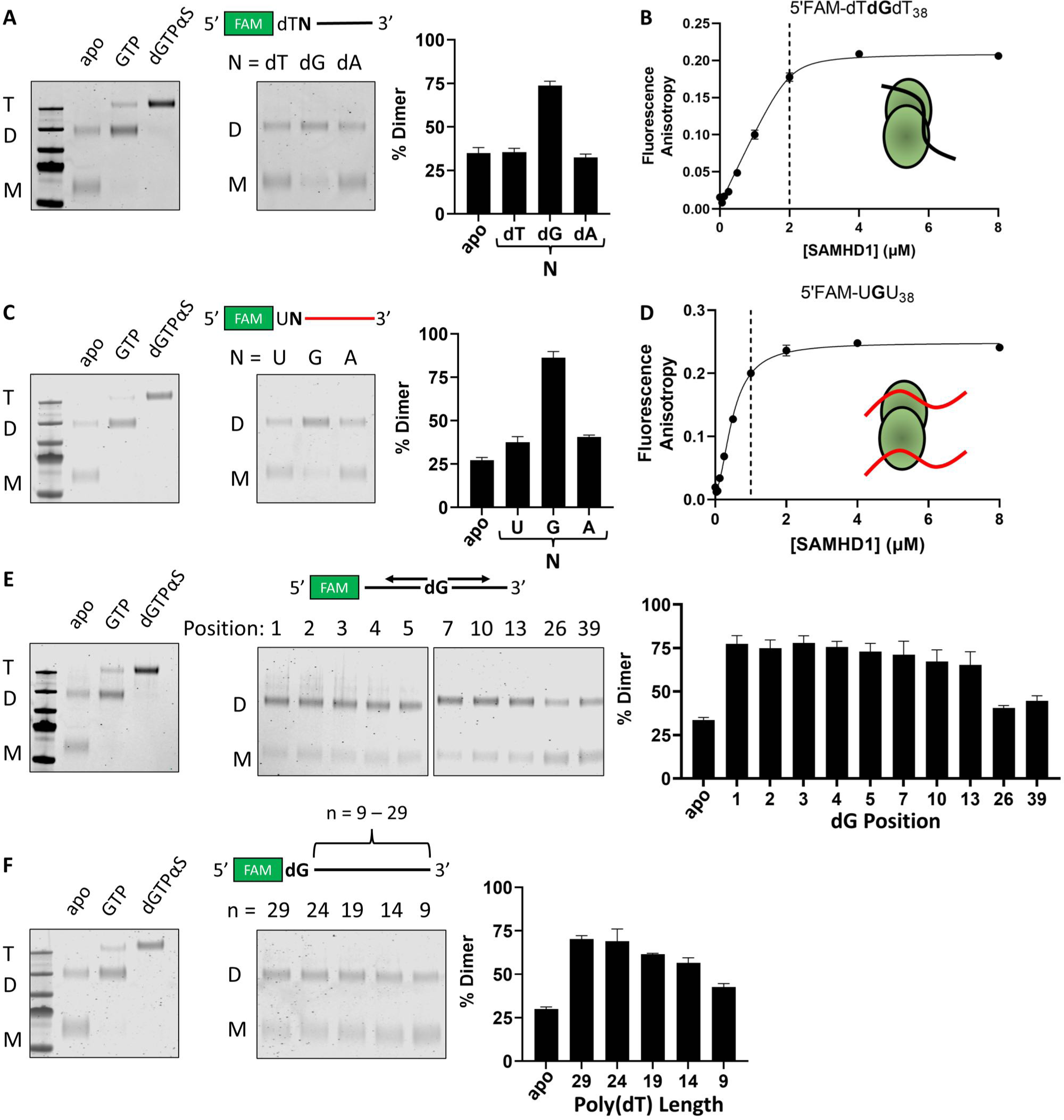
Single G-bases near the 5’ end of ssDNA and ssRNA induce SAMHD1 dimerization. All crosslinking experiments were carried out as follows: SAMHD1 (1 μM) was incubated under each condition for 10 minutes, then crosslinked with 50 mM glutaraldehyde. All gels included dedicated marker lanes for SAMHD1 alone, SAMHD1 and 50 μM GTP, and SAMHD1 and 100 μM dGTPαS to confirm monomer (M), dimer (D) and tetramer (T) species. (**A**) Oligomeric states of SAMHD1 induced by 1 μM 5′FAM-dT**dN**dT38 (where N = dT, dG or dA). (**B**) Binding of SAMHD1 to 5′FAM-dT**dG**dT38 (1 μM) indicates a stoichiometry of two SAMHD1 monomers per DNA strand. (**C**) Oligomeric states induced by 1 μM 5′FAM- U**N**U38 (where N = U, G or A). (**D**) Binding of SAMHD1 to 5′FAM-U**G**U38 (1 μM) indicates a stoichiometry of one SAMHD1 monomer per RNA strand. (**E**) Oligomeric states induced by 5’FAM-labeled dT homopolymer ssDNA 40mers (1 μM) as a function of the position of the dG base within the sequence. (**F**) Oligomeric states induced by dT homopolymers of increasing lengths with a single dG base in position 1 ([DNA] = 1 μM).

In the absence of nucleic acids and in the presence of 50 mM glutaraldehyde crosslinker, 1 μM apo- SAMHD1 is observed as a mixture of 65% monomers and 35% dimers during denaturing polyacrylamide electrophoresis. This increases to ∼85% dimer in the presence of 50 μM GTP, and ∼90% tetramer in the presence of 100 μM dGTPαS (**Fig. 3A**). In the presence of 1 μM 5′FAM-dT**N**T_38_, no observable shift in the ensemble of oligomeric states was observed when N = dT or dA. In contrast, when N = dG the equilibrium was shifted to ∼75% dimer (**Fig. 3A**). Concentration dependent GAXL measurements using 5′FAM- dT**dN**dT_38_ (N = dT, dG) confirmed that no dimerization was induced by 5′FAM-dT_40_ even when it was present at a 10-fold excess over SAMHD1, whereas 5′FAM-dT**dG**dT_38_ induced SAMHD1 dimerization in a concentration-dependent manner (**Supplemental Fig. S6**). Furthermore, these 5′FAM-dT**dG**T_38_-induced dimers form at a 2:1 SAMHD1-ssDNA stoichiometry as indicated by a binding titration using 1 μM [ssDNA](**Fig. 3B**).

Guanosine-dependent dimerization was also observed using 1 μM of the ssRNA 5′FAM-U**N**U_38_ (**Fig. 3C**). Near-homogenous dimer was produced when N = G, whereas no discernable shift in the equilibrium was observed when N = U or A. In contrast to ssDNA, a stoichiometric binding titration revealed that these RNA-induced protein dimers form at a 1:1 SAMHD1-ssRNA stoichiometry (**Fig 3D**). This result indicates that both A1 sites in the RNA-bound dimer are occupied with 5′FAM-U**G**U_38_, whereas only one A1 site of the DNA-bound dimer is occupied by 5′FAM-dT**dN**dT_38_. The underlying basis for the different stoichiometries for binding ssRNA and ssDNA is not currently known.

The dG positional dependence of dimerization was examined by placing the dG specificity residue at ten positions (*n*) from the 5′ end of 5′FAM-dT_N_**dG*^n^***dT_N_-3′ (*n* = 1-5, 7, 10, 13 and 26). For dG in positions 1 to 13, dimerization was found to be efficient and independent of position (**Fig. 3E**). In contrast, a dG at position 26 or 39 was deficient in dimerization. A more limited analysis with ssRNA (5′FAM-U_N_**G*^n^***U_N_-3′ *n* = 2, 13n and 26 revealed the same trend (**Supplemental Fig. S7**), wherein G at position 26 was deficient in dimerization. Furthermore, crosslinking with dT_38_**dG**dT-3’FAM and U_38_**G**U-3’FAM revealed that distance from the 5’FAM label was not the cause of deficient dimerization by 3’-proximate G and dG bases (**Supplemental Fig. S8**). Consistent with this, crosslinking with unlabeled versions of key oligonucleotides from these experiments revealed that 5’ guanine-dependent dimerization is not dependent on the 5’FAM label (**Supplemental Fig. S9**).

The dependence of dimerization on the number of dT nucleotides 3′ to the dG specificity residue was assessed using the constructs 5′FAM-dT**dG**T_9-29_3′ (**Fig. 3F**). Although there was no observed difference in the dimerization efficiency when 29 and 24 nucleotides were present 3′ to the dG, the dimerization dropped by 10%, 15%, and 30% for constructs with only 19, 14 or 9 3′ nucleotides. These effects parallel the length effects on binding affinity and suggest that a stretch of about 20 nucleotides 3′ to the dG residue is needed to fully access the nonspecific DNA binding site (**Supplemental Table S2**).

### Two or more G residues in ssDNA or ssRNA promotes tetramerization of SAMHD1

In contrast to the dimerization observed in the presence of single-G-containing ssDNA and ssRNA, mixed sequence ssDNA and ssRNA (ssDNA_32_ and ssRNA_32_, each containing eight G residues) induced an oligomeric state (T*) with the same electrophoretic mobility as the tetramer (T) induced by dGTPαS (**Fig. 4A, B**). Interestingly, when the concentration of ssDNA_32_ or ssRNA_32_ was equal to or less than the SAMHD1 monomer concentration, homogenous tetramer was observed (**Fig. 4A, B**). In contrast, when the concentration of ssRNA_32_ or ssDNA_32_ exceeded the SAMHD1 concentration, the dimeric state began to predominate at the expense of the tetramer form. From these titrations, we inferred that the apparent tetrameric species (T*) resulted from the association of two SAMHD1 dimers, each bound to one or two ssNA strands. We further surmised that as [ssNA] exceeded [SAMHD1], competing binding modes with the excess NA hindered formation of T*. We observed that ssRNA_32_ binding biased the equilibrium towards tetramer to a greater extent than ssDNA_32_, suggesting that tetramer formation is more favorable with ssRNA than ssDNA. Interestingly, the protein-nucleic acid ratios required to form T* using ssDNA_32_ and ssRNA_32_ were different: ssDNA_32_ formed T* at a 4:1 ratio of protein monomer to DNA strands (**Fig. 4C**) and ssRNA_32_ formed T* at a 2:1 ratio of protein monomer to RNA strands (**Fig. 4D**).

**Figure 4.**
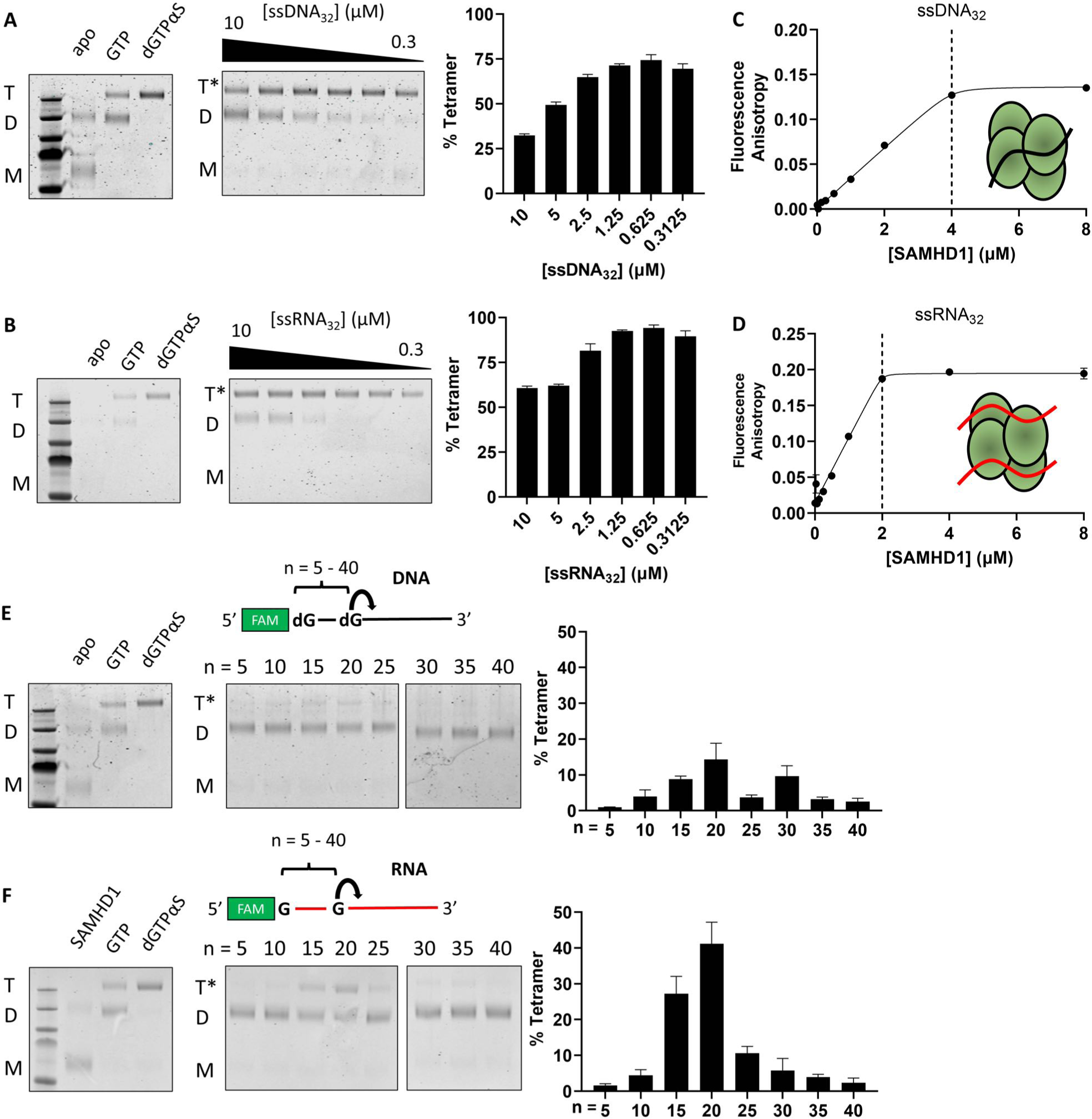
Two or more G bases in ssDNA and ssRNA with ∼20 nt spacing induce SAMHD1 tetramerization. GAXL crosslinking experiments were carried out as in Fig. 3. (**A**) Oligomeric states of SAMHD1 induced by ssDNA32. Lanes from left to right reflect serial two-fold dilutions in the range 10 to 0.3 μM. (**B**) Oligomeric states of SAMHD1 induced by ssRNA32. Lanes from left to right reflect serial two-fold dilutions of each NA in the range 10 to 0.31 μM. (**C**) Stoichiometric binding of SAMHD1 to ssDNA32 (1 μM). (**D**) Stoichiometric binding of SAMHD1 to ssRNA32. (**E**) Oligomeric states induced by a 40mer ssDNA with a single 5′ G base fixed in position 1 and a second G base positioned at the indicated spacings (*n*) (5′FAM- **dG**dTn**dG**dTx). (**F**) Oligomeric states induced by a 40mer ssRNA with a single 5′ G base fixed in position 1 and a second G base positioned at the indicated spacings (*n*) (5′FAM-**G**Un**G**Ux).

Since T* was only observed with ssNA containing multiple G nucleotides, we hypothesized that the baseline requirement for T* formation might be the presence of at least two G residues in the NA strand. To interrogate this hypothesis, we designed a series of 5’FAM-labeled ssRNA and ssDNA 40mers with a single G residue fixed at the 5′ end and a second G residue spaced *n* = 5, 10, 15, 20, 25, 30, 35 or 40 nt from the 5′ end (5′FAM-**G**U_n_**G**U_x_ and 5′FAM-**G**dT_n_**G**dT_x_)(**Fig. 4E, F**). For the ssRNA constructs, there was a distinct spacing (*n* ∼ 20) where tetramerization peaked. With inter-G spacings less than or greater than 20 nucleotides, the amount tetramer was diminished. Thus, T* formation requires two or more G residues spaced roughly twenty bases apart in the context of a ssRNA strand. For the ssDNA constructs, very little T* formation was observed as compared to the mixed sequence ssDNA_32_. The detailed basis for this difference is not known, but it is consistent with the lesser tendency of ssDNA_32_ to form T* as compared to ssRNA_32_.

### Cryo-EM structures of ssRNA32 bound to SAMHD1

To better understand the tetrameric species that formed in the presence of mixed sequence ssNA, we characterized the ssRNA_32_-induced tetramer (T*) using cryogenic electron microscopy (cryo-EM). Collection of 26,000 dose-fractionated movies of SAMHD1 (6.2 μM) mixed with ssRNA_32_ (3.5 μM) produced heterogeneous mixture of RNA-bound complexes consisting of dimers (40% of particles picked), tetramers (40% of particles picked), and a small number of hexamers (20% of particles picked) (**Supplemental Fig. S10**). The basis for the greater cryo-EM particle heterogeneity as compared to the crosslinking results is not known, but may arise from loose complexes dissociating during the vitrification process^23^. Nevertheless, the presence of dimers and tetramers—which are also observed in our extensive biochemical studies—suggested a pathway for formation of T* (**Fig. 5A**). Rigid body docking of SAMHD1 dimers into the observed particles suggests that the first step on the pathway to T* involves the association of two ssRNA-bound dimers (D) to form a D•D complex (**Fig. 5A**). Although RNA is not directly observed in the D or D•D complexes due to the conformational heterogeneity, its presence is inferred based on the biochemical measurements above. The transient D•D complex then undergoes a conformational rearrangement to form more compact tetrameric complexes that are in dynamic equilibrium (T*_op_ and T*_cl_, refined to 3.04 Å and 3.44 Å respectively) (**Fig. 5A**). The most compact conformation, T*_cl_ (right), showed continuous density for two bound RNA strands (red, **Fig. 5A**), each bridging two A1 sites of different dimers. In contrast, the RNA density for the more open conformation T*_op_ (left) was discontinuous as the RNA strand moved away from either A1 site. Due to its superior RNA map quality, our remaining discussion focuses on conformation T*_cl_. The hexameric particles that were observed could not be further studied due to their low population in the dataset and strong orientational preference (**Supplemental Figure S10**). The biological significance of the hexamer (if any) is not known. However, we note that a hexameric form of a bacterial homolog of SAMHD1 has been reported^24^, and higher-order SAMHD1-nucleic acid complexes have been previously predicted by live cell imaging experiments^25^.

**Figure 5.**
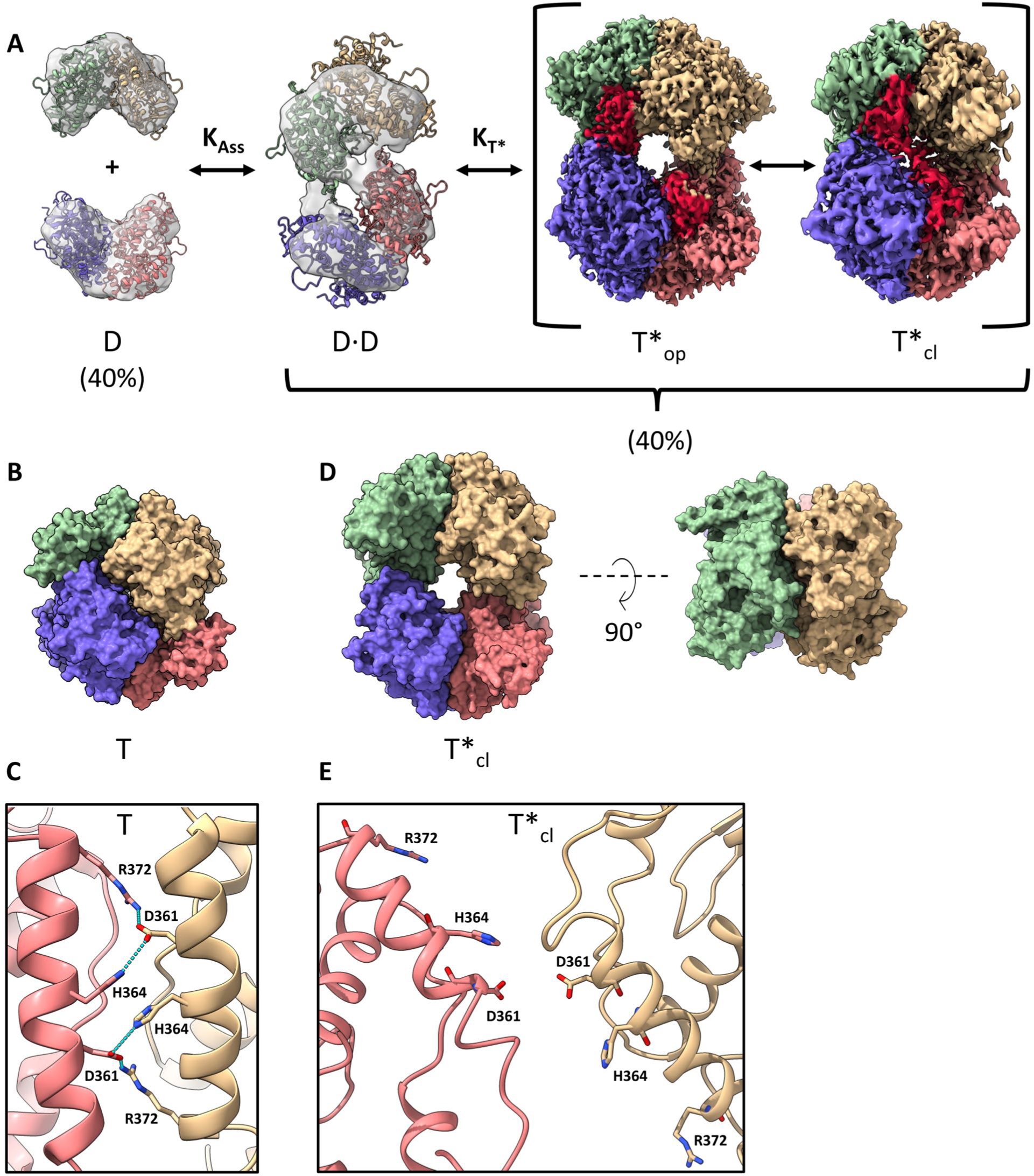
CryoEM analysis of ssRNA32 complexes with SAMHD1. (**A**) Overview of particles observed in cryo-EM study and proposed mechanism of T* formation. Abbreviations: D (dimer), T*op, T*cl (open and closed conformational states of the ssRNA bound tetramer). For low resolution species, (D and D·D) SAMHD1 dimers were rigidly docked into the density using the fitmap command in ChimeraX. For both conformations of T*, density corresponding to RNA is colored violet. (**B**) Structure of the canonical dNTP-saturated tetramer induced by dGTPαS (T, PDB: 7UJN). (**C**) Dimer-dimer interface interactions in the T complex with dGTPαS. A charged hydrogen bonding network involving the interfacial residues D361, H364, and R372 is observed. (**D**) Structure of the ssRNA32-bound tetramer (T*cl). The RNA was deleted from this depiction of T*cl to facilitate comparisons between T and T*cl. (**E**) Dimer-dimer interface interactions in the T*cl. complex with ssRNA32. A significant displacement of the two interfacial helices disrupts the hydrogen bonding of residues D361, H364, and R372.

There are significant differences in the packing of SAMHD1 dimers between the canonical nucleotide-bound tetramer (T) and the T*_cl_ complex (**Fig 5B**, **D**). Upon formation of T*_cl_, only 983 Å^2^ of solvent exposed surface area is buried compared to 6762 Å^2^ for T, and numerous side chain interactions form along the dimer-dimer interface of T that are absent in the T*_cl_ complex. One example is the hydrogen bonding network formed by helical residues R372, H364, and D361 (**Fig. 5C**), which are splayed apart in T*_cl_ (**Fig 5E**). Previous mutations of R372 and H364 have indicated their role in dNTPase activity and tetramer stability^5^.

We were also curious whether RNA density might be observed in the HD-domain active site. Using our previous cryo-EM structure of the dGTPαS-bound T complex for comparison, we found that that the T*_cl_ active site was devoid of any nucleotide density (**Supplemental Figure S11A**, **B**), more compressed, and partially occluded from solvent by a random coil peptide chain consisting of residues 503-510 (**Supplemental Figure S11C, D**). Thus, the T*_cl_ structure suggests that the active site is not easily accessible by dNTPs or RNA nucleotides. Consistent with the structural observations, activity measurements on T* did not detect any dNTPase or RNase activity (see below). The lack of activity cannot be attributed to low occupancy of the catalytic iron metal because density for the metal is observed in T*_cl_ and the measured stoichiometry for enzyme bound iron is 0.96 iron atoms per enzyme monomer (ICP-MS) **(Supplemental Figure S11A, B)**.

Lacking the network of protein-protein interactions that ordinarily stabilize SAMHD1 tetramers, T*_cl_ is instead stabilized by a novel, RNA-mediated tethering mechanism (**Fig. 6A**). Gaussian smoothing of the density map reveals a continuous tube of RNA density bridging the A1 sites of two dimers (i.e., each RNA strand bridges two A1 sites). Although high-resolution refinement of the flexible regions of the RNA sequence was not possible, molecular dynamics flexible fitting was used to obtain a reasonable pose for the bound RNA (**Fig. 6B**)^26^. This procedure indicated that residues G8 and G25 in ssRNA_32_ can be positioned in the A1 sites and fit the observed tube of density in T*_cl_. However, G8-G25 is one of three pairs of guanine residues in the sequence that can fit the density and satisfy the ∼20 nt spacing requirement for tetramerization (G8-G24, G8-G25, G8-G30). Importantly, these ambiguities in the binding register do not alter the major finding that the RNA strand bridges two A1 sites.

**Figure 6.**
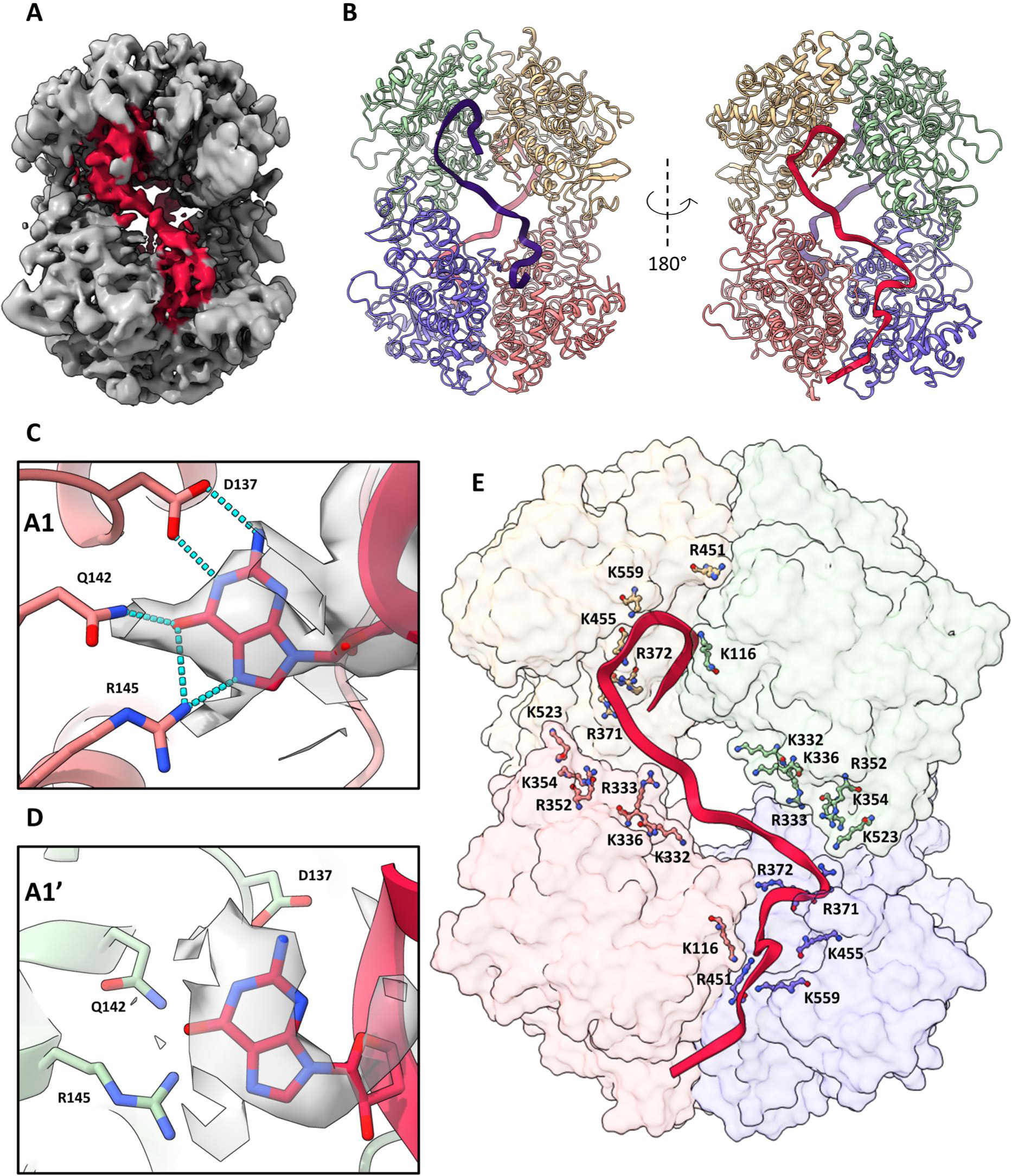
Molecular basis for ssRNA32 binding (T*cl). (**A**) Smoothed density map of T*cl shown at low contour to highlight ssRNA connectivity between allosteric sites. Density corresponding to ssRNA32 is highlighted in pink. (**B**) Fragments of ssRNA32 can be posed in the tube density through molecular dynamics flexible fitting using ISOLDE positioning G8 and G25 in the A1 sites on each face of the tetramer. (**C**) A guanosine nucleotide modeled in the A1 site of chains A (purple) & B (pink), where it forms a hydrogen bond network with D137, Q142, and R145. (**D**) A guanosine nucleotide modeled in the A1 sites of chains C (green) and D (yellow) (denoted A1′). The guanosine binding pose is rotated as compared to panel (**C**) and does not form hydrogen bonds with D137, Q142, and R145. This difference in poses between A1 and A1′ sites may arise from the different 5’-3’ orientation of the RNA strand at the A1′ sites. (**E**) Cationic residues located near the dimer-dimer interface of T*. Residues K116, K332, R333, K336, R352, K354, R371, R372, K377, R451, K455, K523, K559 are positioned to interact with the flexible sugar-phosphate backbone of ssRNA32.

The high-resolution (lower) dimer of T*_cl_ shows unambiguous A1 site density for a guanine base (G8), which forms the same network of hydrogen bonds with residues D137, Q142, and R145 observed with the GTP complex (**Fig. 6C**). In addition, the densities flanking the guanine were consistent with the sequence containing G8 (5’C_7_G_8_G_9_3’). In contrast, the lower resolution (upper) dimer of T*_cl_ shows less well-defined density for the guanine nucleotide (G25) in its A1’ site (**Fig. 6D**), which may be attributed to the loop in the RNA chain required to access the site in the correct orientation (**Fig. 6E**). Due to the lesser RNA map quality of the upper dimer, the exact orientation of this loop is unclear. Nevertheless, symmetric A1 site binding by G residues within a single ssRNA chain is required to maintain the same 5’- 3’ binding polarity at both sites. From these data, we cannot exclude the possibility that different pairs of guanines may dynamically bind two A1 sites. This may explain the less well-defined T*_op_ conformation, as well as other more transient tetrameric species in the dataset. In addition to guanine-specific binding at the allosteric sites, T*_cl_ also possesses numerous cationic residues near the flexible RNA strands which are well-positioned to interact non-specifically with the sugar-phosphate backbone (**Fig. 6E**). Several of these residues have been previously implicated in RNA binding by mutagenesis^5^.

### RNA-bound monomer, dimer and tetramer forms of SAMHD1 do not have dNTPase or RNase activity

Because NTP-induced tetramerization (T) is the canonical activation mechanism of SAMHD1, we tested whether T* formed in the presence of ssRNA_32_ might also stimulate SAMHD1 dNTPase activity. For this assessment, we used our validated 2-[^14^C]-dTTP thin layer chromatography assay for dTTP hydrolysis (**Supplemental Figure S12**). As expected, 0.5 mM GTP stimulated hydrolysis of 1 mM dTTP, and no dTTP hydrolysis was observed in the absence of GTP (**Supplemental Figure S12A**). In the presence of 0.25 to 2 μM ssRNA_32_ and 0.5 μM SAMHD1, we detected no dTTP hydrolysis over the same 30 min time frame (**Supplemental Figure S12B, C**). Consistent with the structural observations, the activity measurements indicate that binding of ssRNA_32_ does not lead to a productive conformation of the active site.

Given the recent report that SAMHD1 RNA exonuclease activity was involved in cellular RNA homeostasis^15^, we investigated whether ssRNA-induced dimerization and tetramerization might regulate RNase activity. Three synthetic ssRNAs were used as potential substrates (i) a polyU 40mer (5′FAM-U_40_) that binds to monomeric SAMHD1 and does not promote oligomerization, (ii) polyU with a single 5′ guanine (5′FAM-U**G**U_38_), which induces dimeric SAMHD1, and (iii) 5′FAM-ssRNA_32_ that induces tetrameric SAMHD1. Incubation of these substrates (1 μM) with SAMHD1 (1 μM) for two hours resulted in no more than 5% degradation of the substrate (**Supplemental Figure S13 A-D**). We conclude that stringently purified SAMHD1 does not possess RNase activity with a variety of synthetic ssRNA substrates, confirming our previous findings (**Supplemental Figure S14**)^6^. We previously concluded that trace nuclease contamination in SAMHD1 protein isolates was correlated with the observed low level of RNase activity. Consistent with this interpretation, we performed a mass spectrometry proteomics analysis of SAMHD1 fractions after the Ni-NTA, SP-sepharose and size exclusion chromatography steps and detected trace levels of three ribonucleases (*E. coli* polynucleotide phosphorylase, ribonuclease E, and RpsP) and a DNase (uvrA) that were removed or greatly diminished over the course of the purification. These data and supporting controls are provided in **Supplemental Table S4** and **Supplemental Figure S15**.

### SAMHD1 forms “beads on a string” complexes with long ssRNA

The structural studies with ssRNA_32_ suggested that longer RNA strands with multiple guanine bases at random spacings might result in single strands of RNA looping around dimeric or tetrameric SAMHD1 with guanines occupying two or four A1 sites for dimer and tetramer species, respectively. To explore this further, we *in vitro* transcribed a 2.1 kb RNA using recombinant T7 RNA polymerase and collected negative stain transmission electron micrographs of SAMHD1 bound to the RNA (**Supplemental Figure S16**)(**Fig. 7**). Micrographs of the ssRNA alone showed a highly collapsed molecule (**Fig. 7A**). In contrast, addition of SAMHD1 led to extended RNA structures with punctate globular particles distributed along the length of the chain (**Fig. 7A**). As expected from the biochemical and cryo-EM results, the sizes of the particles were consistent with both dimers and tetramers of SAMHD1. Identical micrographs were obtained of T* complexes with ssRNA_32_, which as expected, only showed isolated, mostly tetrameric particles (**Fig. 7B**). Based on all of the structural and biochemical findings we propose a model where SAMHD1 binding to long ssRNA molecules leads to the formation of enzyme dimers and tetramers that are in dynamic equilibrium depending on the protein binding density and RNA sequence (**Fig. 7C**).

**Figure 7.**
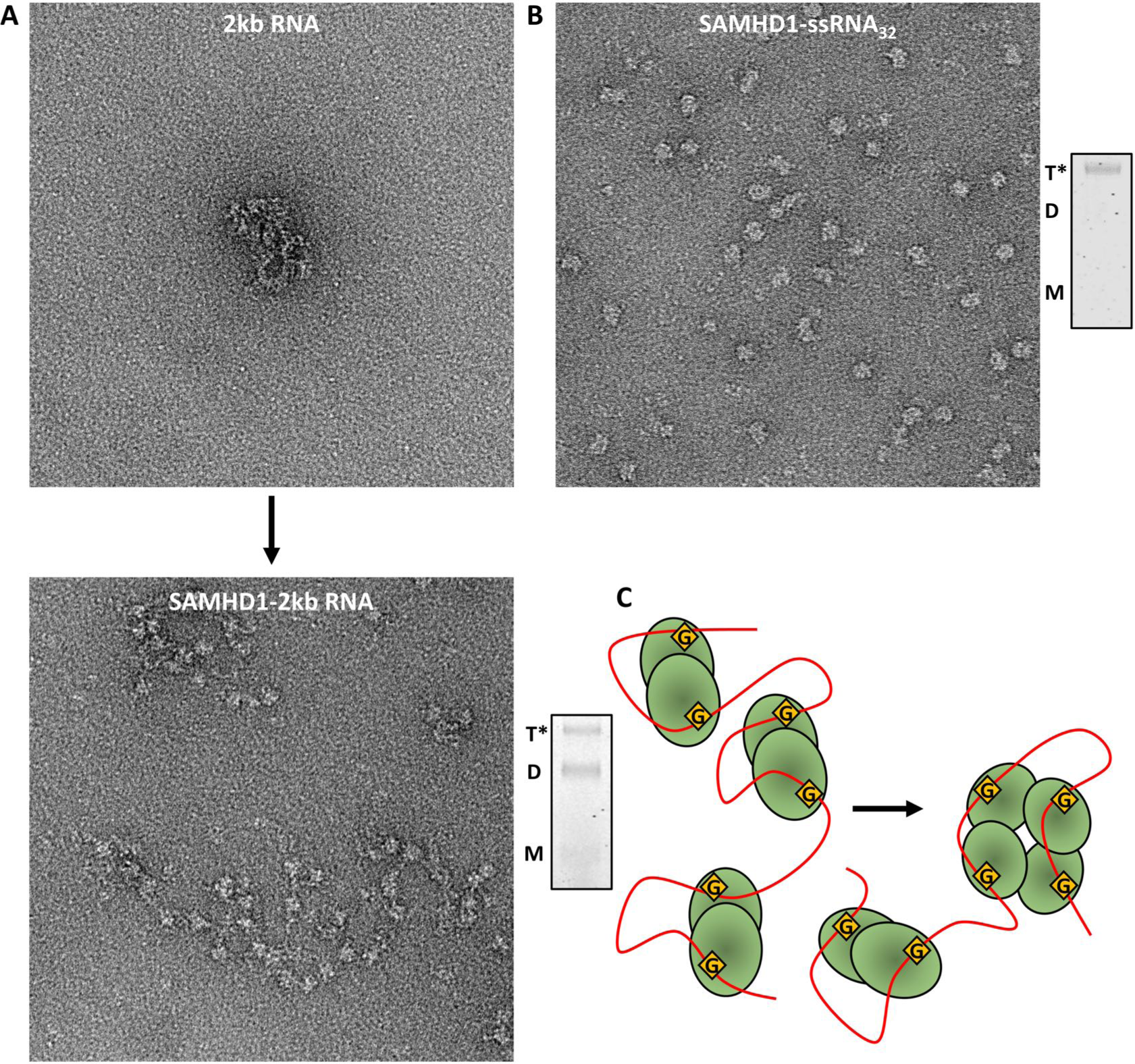
SAMHD1 interacts with long mRNA molecules as an ensemble of dimers and tetramers. Transmission electron micrographs (TEM) shown at 150,000X magnification using a 1% uranyl formate negative stain. (**A**) TEM image of in vitro transcribed 2 kb RNA (10 ng/μL) alone (top) and in the presence of 100 nM SAMHD1 (bottom). The mixed dimeric (D) and tetrameric (T*) states of SAMHD1 observed in the images were confirmed by the GAXL method (gel lane on the right). (**B**) SAMHD1 (100 nM) bound to ssRNA32 (100 nM). The largely tetrameric (T*) state of SAMHD1 observed in the images was confirmed by the GAXL method (gel lane on the right). (**C**) Proposed model for SAMHD1 binding to long RNA involves G binding to A1 sites and the formation of enzyme dimers. Where the spacing of guanine residues is correct, dimeric units can coalesce to form tetramer.

## DISCUSSION

Our results begin to unravel how the *in vitro* nucleic acid binding of SAMHD1 relates to its known activities in cells. First, specificity for ssNA binding relies on recognition of G residues that are exposed in single-stranded DNA and RNA. Binding of guanine residues to the A1 site serves to destabilize the dNTPase tetramer by displacing GTP^7^, and then tethers a single NA strand to multiple A1 sites if more than one guanine with appropriate spacing is present (**Fig. 7C**). Thermodynamically, there is potential for a reduced entropic cost for binding of multiple G bases tethered in a flexible NA strand as compared to binding of free GTP. Such effects could allow ssNA to compete effectively with GTP through substantial avidity effects. Although our structural studies focus on ssRNA, we speculate that a similar binding mechanism may apply to duplex DNA with 5′ single strand regions that contain guanines. Such DNA structures are associated with replication forks, double strand breaks, and R loops and SAMHD1 is known to associate with these structures in cells ^8–10, 12, 13^.

As of yet, the full biological significance of the unique guanine-centric nucleic acid binding mode of SAMHD1 remains unclear. Indeed, to our knowledge this is the first reported example of a GTP binding site being repurposed for NA binding. As already mentioned, the requirement for exposed guanines discriminates against binding to duplex structures, but it may also contribute to the reported localization of SAMHD1 to guanine enriched regions of the genome such as telomeres^14^. Although guanine is depleted in some genomic DNA regions, guanine bases are generally present at sufficient densities in the genome that the binding modes observed here would be widely available.

Our TEM images strongly suggest that binding of SAMHD1 can disrupt the secondary structure present in most ssRNA molecules (**Fig. 7**). This observation is intriguing given the recent report by Maharana *et. al.* implicating SAMHD1 in modulation of RNA liquid condensates in the cytosol and its suppression of MDA5-dependent autoimmunity^15^. Since many viral RNA sensors specifically target double stranded RNA, our findings highlight a possible ssRNA-mediated, immunosuppressive role of SAMHD1. Of note, ADAR1, another gene product implicated in Aicardi-Goutières Syndrome, also suppresses autoimmunity by destabilizing double-stranded RNA structures^27^. Given the well-established role of SAM domains in protein-protein interactions^28^, it is easy to envision SAMHD1 as an RNA scaffold protein to recruit additional factors (such as an exonuclease) required for RNA homeostasis.

The present structural and RNase activity measurements do not provide support for a putative exonuclease activity of SAMHD1. While ssRNA is observed in the allosteric sites and tetramer interface, no RNA density is found in the HD-domain active site, nor does the RNA binding mode direct the 3′ RNA ends towards the active site. Furthermore, the observed beads-on-a-string binding mode of SAMHD1 on long ssRNA strands is not consistent with 3′-5′ exonuclease activity, and in fact, would sterically hinder such a process. The trace proteins detected in our proteomics analysis suggest that RNase contamination is possible, especially in some previous studies where single-step purifications were used. It remains to future structural studies how SAMHD1 interacts with its established partners in DNA repair (MRE11, RPA, and CTIP), as well as putative partners in RNA homeostasis.

### Methods Chemicals

2’-Deoxythymidine-5’-triphosphate (dTTP) was obtained from Promega. 2-^14^C labeled 2’- deoxythymidine-5’-triphosphate (2-^14^C-dTTP) was obtained from Moravek biochemicals. Guanosine-5’- triphosphate (GTP) was obtained from ThermoFisher Scientific. Xanthosine-5’-triphosphate (XTP) and glutaraldehyde were obtained from Sigma-Aldrich. 2’-deoxyguanosine-5’-[α-thio]-triphosphate (dGTPαS) was obtained from TriLink Biotechnologies. Uranyl Formate was obtained from Electron Microscopy Sciences. C18 reversed-phase thin layer chromatography (TLC) plates were obtained from Macherey-Nagel.

### DNA and RNA oligonucleotides

All oligonucleotides were synthesized by Integrated DNA Technologies (IDT). DNA oligonucleotides greater than 40 bases and all RNA oligonucleotides were HPLC purified. All oligonucleotides dissolved to a 100 μM concentration in nuclease-free water. Sequences are listed in **Supplemental Table S1**.

### *In vitro* transcription of 2kb RNA

RNA encoding the SAMHD1 gene was transcribed from a pET19b plasmid with 10X-His SAMHD1 under a T7 promoter HiScribe T7 High Yield RNA transcription kit (NEB). The resulting RNA was purified using a Monarch RNA cleanup kit (NEB), aliquoted, and flash frozen in liquid nitrogen.

### Protein expression and purification

Human SAMHD1 wild-type or D137N harbored in a pET19b plasmid as a PreScission protease cleavable 10xHis fusion construct was expressed in chemically competent BL21(DE3) E. coli cells (Agilent). An overnight starter culture of cells was grown in 2xYT supplemented with carbenicillin (50 µg/L) and subsequently inoculated 1:100 v/v in 2xYT media. Cultures were grown in a Harbinger LEX-48 parallel bioreactor system at 37 °C until an OD_600_ of 0.7 was reached. The cells were then cold-shocked for 30 minutes in an ice bath, induced with 1 mM isopropyl β-D-1-thiogalactopyranoside (ThermoFisher), and incubated for 20 hours at 20 °C. Cells were harvested by centrifugation (10,000 x g) and stored at −80 °C.

Cell pellets were thawed and resuspended in lysis buffer (50 mM HEPES—pH 7.5, 300 mM KCl, 4 mM MgCl_2_, 0.5 mM TCEP, 25 mM imidazole, and 10% glycerol) with one tablet of protease inhibitor cocktail (Pierce), 1 mg DNase I (Roche), 1 mg RNAse A (Alfa-Aesar), and 5 mg lysozyme (*Amresco*) per 50 mL of buffer. The resuspension was passed two times through a LM10 microfluidizer (Microfluidics) and centrifuged at 40,000 x g to produce a clarified lysate. The lysate was loaded onto a charged, pre-equilibrated 10 mL nickel column (HisPur Ni-NTA resin from ThermoFisher). The loaded column was given stringent, incremental washes with 30, 40, and 50 mM imidazole. After the UV trace returned to baseline, SAMHD1 was eluted from the column using 300 mM imidazole.

SAMHD1 isolates were dialyzed against 4 L of general buffer (50 mM HEPES—pH 7.5, 300 mM KCl, 4 mM MgCl_2_, 0.5 mM TCEP, and 10% glycerol) overnight with 1 mg GST-tagged PreScission protease added to remove the imidazole and 10xHis tag. The following day, the isolate was gently stirred at 4 °C with 2 mL glutathione-agarose resin, then centrifuged to pellet the resin and removed GST-PreScission protease. The SAMHD1 solution was concentrated to ∼10 mg/ml, then diluted 16-fold in S-column binding buffer (50 mM HEPES—pH 7.5, 20 mM KCl, 4 mM MgCl_2_, 0.5 mM TCEP, and 10% glycerol) and applied to a 10 mL GE HiTrap SP sepharose fast-flow column (two 5 mL cartridges in tandem) at 0.5 mL/minute. SAMHD1 was eluted using an 80 ml gradient from 20 to 500 mM KCl. The SAMHD1 isolate was again concentrated to ∼10 mg/ml and injected 45 mgs at a time onto a Cytiva Superdex 200 pg HiLoad 26/600 size exclusion column to remove aggregates. The resulting SAMHD1 isolate was concentrated to 8-9 mg/ml and flash frozen in liquid nitrogen in 30 μL aliquots.

### Metal content of purified SAMHD1

Purified SAMHD1 protein (∼3 mg in 400 μL storage buffer) was subjected to a 6-hour dialysis against 600 mL of buffer (30 mM HEPES—pH 7.5; 230 mM KCl, 3 mM MgCl₂ and 0.5 mM TCEP) using a 10 kDa cut- off dialysis chamber. The buffer had been pre-treated with Chelex resin to remove trace metals. The sample was then dialyzed overnight against the same buffer containing 10 g/L Chelex resin. The dialyzed protein was filtered through a 0.2 micron membrane and ICP-MS measurements were performed at the University of Massachusetts Amherst, Mass Spectrometry Facility using Perkin-Elmer NexION 350D ICP-MS instrument. Briefly, the protein sample (171.2 μM) and identically treated buffer controls were treated with trace metal nitric acid (50 µL of sample was added to 1.95 mL of 5% HNO₃), vortexed, and centrifuged (5000×g, 5 min). The clear supernatants were diluted (1:20) and used for ICP-MS analyses. Linearity of detection for iron was assessed using an ⁵⁷Fe calibration standard at nine concentrations ranging from 0–200 ppb. A stoichiometry for enzyme bound iron was calculated from the measured iron concentration in the sample and the calculated protein concentration based on its extinction coefficient at 280 nm (80604.1 M^-1^ cm^-1^).

### Proteomic Analysis of purified SAMHD1

One-hundred and fifty micrograms of protein from each of the three SAMHD1 purification steps (post-Ni affinity, post-S Sepharose and post-SEC) were analyzed as described in **Supplemental Method S2**.

### Fluorescence anisotropy measurements of ssNA binding

The fluorescence anisotropy measurements were carried out in Corning Costar black, flat-bottom, 96- well assay plates using an Agilent BioTek Synergy Neo2 plate reader maintained at 25°C. Binding reactions mixtures were prepared in binding buffer (50 mM HEPES—pH 7.5, 50 mM KCl, 1 mM EDTA, 0.5 mM TCEP) with 50 nM FAM-labeled oligonucleotide and allocated in the assay plate (70 μL/well) using a multichannel pipet. SAMHD1 (8 μM) was added to the first column and then serially diluted down the plate. Three replicate titrations were performed for each oligonucleotide. Polarization of the light emitted from the FAM fluorophore was measured using a 485/525 nm polarization filter cube. Polarization was plotted as a function of SAMHD1 concentration and converted to fluorescence anisotropy using the following formula:

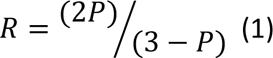

Where P is fluorescence polarization at a given point in the titration and R is fluorescence anisotropy at a given point in the titration. The resulting anisotropy vs. total [SAMHD1] curves were fit to a Hill binding equation:

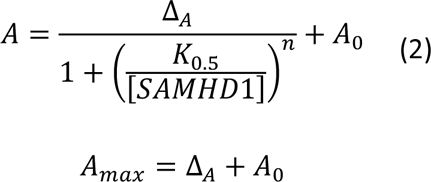

where R is the observed anisotropy, A_0_ is the baseline anisotropy of the free oligonucleotide, Δ_A_ is the change in anisotropy over the course of the titration, A_max_ is the anisotropy at the end of the titration, [SAMHD1] is the total SAMHD1 concentration, K_0.5_ is the [SAMHD1] which produced half-maximal change in anisotropy, and n is the cooperativity factor.

### Competition displacement experiments

Competition displacement experiments were performed in the same manner as standard binding measurements, with the exception that unlabeled competitor was added and serially diluted in a binding mixture containing 1 μM SAMHD1 and 0.5 μM FAM-labeled ssNA. These complex equilibria were modeled using the DynaFit numerical integration program (**Supplemental Method 1**).

### Stoichiometric binding experiments

Stoichiometric binding experiments were performed in the same manner as standard binding measurements, except they were carried out at 1 μM ssNA and analyzed using a quadratic binding equation as previously described^7^.

### SAMHD1 dNTPase activity

SAMHD1 (0.5 μM) was incubated with dTTP (1 mM) and GTP or XTP (0.5 mM) for 32 minutes in assay buffer (50 mM HEPES–pH 7.5, 50 mM KCl, 5 mM MgCl_2_, 0.5 mM TCEP) with 20 nCi [2-14C]-dTTP labeling. Three replicate reactions were carried out under each condition with SAMHD1 as the initiator. One-microliter fractions were withdrawn at 0, 0.5, 1, 2, 4, 8, 16, and 32 minutes and quenched by spotting onto a C18-reversed phase TLC plate. TLC plates were developed in 50 mM KH_2_PO_4_ (pH 4.0) to separate the substrate dTTP from the product dT. Developed TLC plates were exposed on a GE storage phosphor screen overnight and scanned on a Typhoon Imager (GE Healthcare). Substrate and product signal was quantified using ImageJ. The amount of product formed at each time point in each replicate was calculated using eq 3:

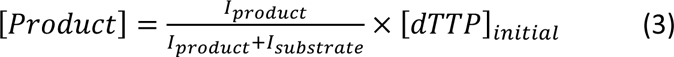

where I_product_ is the signal intensity of the dT product peak, I_substrate_ is the signal intensity of the dTTP substrate peak, and [dTTP]_initial_ is the initial concentration of dTTP substrate in the reaction. Initial rates of product formation were obtained from plots of [dT] vs. time, with rates corresponding to slope, and error corresponding to standard error of the slope as determined by linear regression analysis.

### SAMHD1 ribonuclease activity

SAMHD1 (1 μM) was incubated with 5’FAM-U_40_, 5’FAM-UG_40_, or 5’FAM-ssRNA_32_ (1 uM) for two hours in assay buffer (50 mM HEPES–pH 7.5, 50 mM KCl, 5 mM MgCl_2_, 0.5 mM TCEP). Five-microliter fractions were withdrawn at 5, 10, 15, 30, 60, and 120 minutes and quenched in 5 μL formamide loading buffer (98% formamide, 10 mM EDTA). Positive and negative controls were obtained by two-hour incubation with 1 μM bovine pancreatic ribonuclease A (ThermoFisher Scientific) or by adding SAMHD1 storage buffer into a mock reaction. Ladder was made by making an equimolar mixture of the 5’ FAM-labeled ssDNA 30mer, 25mer, 20mer, 15mer, and 10mer used for the length dependence experiment in **Fig. 3b**. All samples were analyzed using an 8M urea, 20% 19:1 acrylamide:bisacrylamide TBE gel run at 25 W for 75 minutes or until the yellow marker dye (TriLink Marker Dye, ThermoFisher Scientific) reached the bottom of the gel.

### Oligomerization measurements using glutaraldehyde crosslinking

SAMHD1 (1 μM) was incubated with GTP (50 μM), dGTPαS (100 μM), or nucleic acids (1 μM) for 5 minutes in binding buffer (50 mM HEPES–pH 7.5, 50 mM KCl, 1 mM EDTA, 0.5 mM TCEP), then crosslinked in 50 mM glutaraldehyde for 10 minutes. For the dGTPαS tetramerization control, 1 mM EDTA was omitted and 5 mM MgCl_2_ was included. Crosslinking was halted by the addition of 170 mM Tris–pH 7.5. Crosslinked fractions were loaded onto a 1.5 mm, 10-well, 4-12% acrylamide gradient Bis-Tris gel (Invitrogen) and ran at 200 V for 40 minutes to separate tetrameric, dimeric, and monomeric species. Gels were stained with Coomassie R250 dye. A PAGEruler pre-stained molecular weight ladder (Thermo) was used to identify which bands corresponded to the different oligomeric states of SAMHD1. Three replicate reactions were performed for each condition.

### Negative stain transmission electron microscopy (TEM) of ssRNA-bound SAMHD1

SAMHD1 (100 nM) was mixed with ssRNA in binding buffer (50 mM HEPES–pH 7.5, 50 mM KCl, 1 mM EDTA, 0.5 mM TCEP), then spotted onto a glow-discharged 300 mesh carbon-film coated copper electron microscopy grid (Electron Microscopy Sciences). Once applied, excess protein sample was blotted away with filter paper. The grid was stained in 1% uranyl formate solution, washed with water, blotted with filter paper, and left to dry for 30 minutes in a dark room. Electron micrographs were taken in Velox at 150,000X magnification on a ThermoFisher Talos L120C TEM equipped with a Ceta camera.

### Cryo-EM of ssRNA-bound SAMHD1

SAMHD1 (6.2 μM) and ssRNA_32_ (3.5 μM) were mixed in dilution buffer (50 mM HEPES—pH7.5, 50 mM KCl, 1 mM EDTA, 0.5 mM TCEP) and incubated for 10 minutes at room temperature. The SAMHD1-ssRNA32 complex was then applied to a plasma cleaned (ArO_2_) Quantifoil R2/1 200 mesh gold grid and vitrified in liquid ethane (Vitrobot Mark IV, ThermoFisher Scientific). Grids were imaged using a Titan Krios G2 transmission electron microscope operated at 300 kV, equipped with a Falcon 4i direct electron detector and Selectris X imaging filter. All screening and data collection procedures were conducted in EPU (Thermo Fisher Scientific). Movies in EER format were obtained at a magnification of 165,000x (a pixel size of 0.733 Å), with a total electron dose of 40 e-/Å2. A total of 26,702 movies were collected. The raw frame stacks were subjected to gain normalization, alignment, and dose compensation using Motioncor2^29^ with patch-based alignment (5 × 5) and without binning. CTF parameters were estimated from the aligned frame sums using CTFFIND4.1^30^.

Motion-corrected images were imported to cryoSPARC 4.0^31^and automated particle picking was performed on a dataset of 4,000 selected images using Blob Picker, followed by 2D classification to generate 2D references. These references were then employed in the template picker for the entire set of micrographs. Blob-based autopicking was applied to the complete set of micrographs. The resulting particles from both the blob picker and template picker were merged, and redundant particles within a 20 Å distance were eliminated. The initial dataset consisted of 92,706,714 particles extracted at 2.93 Å/pixel. Four rounds of 2D classifications were performed on these particles. During the classification process, variable oligomeric status was observed, prompting further 2D classifications targeting specific conformations. Three distinct particle groups were identified: dimer, tetrameric, and hexameric particles. There were additional particles whose oligomeric status could not be reliably validated. A subset of the dimeric particles underwent additional 2D classifications followed by homogeneous and non-uniform refinement for 3D reconstruction. The resulting map showed a good fit with the dimeric SAMHD1 structure (‘D’ in **Fig. 5A**). A subset of 571,935 particles, which exhibited a tetrameric status, was utilized for the generation of an initial model used in the subsequent 3D classification. 2,653,093 particles selected from four rounds of 2D classifications underwent extensive 3D classifications using Relion 4.0^32^ to address data heterogeneity. Classes with optimal secondary structural features were further refined through additional 3D classifications. One of the classes from the first 3D classification revealed a loosely formed tetramer (‘D·D’ in **Fig. 5A**). Three rounds of 3D classifications with incremental increases in regularization (T = 4, 6, 8) were conducted. The third 3D classification revealed two types of tetramers: loose tetramers and tighter tetramers. The latter exhibited additional density at allosteric sites. Particles from the selected class were re-extracted at 1.47 Å/pix and subjected to further 3D classification with a regularization parameter of T=10. The results unveiled two major classes, referred to as T*_op_ and T*_cl_. The particle data was exported to cryoSPARC, where ab-initio modeling, homogeneous refinement, non-uniform refinement, and local refinement were carried out. Final maps were subjected to map-modification implemented in Phenix with two independent half maps and corresponding mask and model as input.

### Model building and refinement

An atomic model of full-length SAMHD1 was generated using AlphaFold^33^ and rigid-body fitted into the cryo-EM maps of T*_op_ and T*_cl_. The model was then iteratively built into the map using Coot (v0.9.8.)^34^in conjunction with real-space refinement in PHENIX^35^. To improve the model quality, residues in loop regions, RNA residues, and side chains with weak or ambiguous density were removed. This iterative process resulted in the production of the final refined models. The complete cryo-EM data processing workflow and validation metrics can be found in **Supplemental Figure S10**. Renderings in this figure were generated UCSF Chimera version 1.16^36, 37^. Renderings in **Figure 5** and **Figure 6** were generated in ChimeraX version 1.5^36, 37^. Statistics pertaining to the model refinement can be found in **Supplemental Table S3**.

### Molecular Dynamics Flexible Fitting with ssRNA32

Molecular dynamics flexible fitting was carried out in ChimeraX version 1.5 using ISOLDE. The ssRNA_32_ sequence was built using the “rna” command and dragged into place in T*_cl_ under global simulation in an AMBER forcefield with a map weighting of 0.45. Residues G8 and G25 in the ssRNA_32_ sequence were positioned in A1 and A1’ respectively on each side of T*_cl_, then subjected to position restraint (spring constant = 50 kJ/mol·Å^2^) while the remaining RNA sequence was positioned in the tube of density bridging the two sites using localized simulations about individual RNA residues. The flexible unbound 5’ and 3’ ends of the RNA were truncated to the ends of the observed density in the T*_cl_ map. The purple and red strands in **Figure 6B** correspond to residues 5-27 and residues 1-27 respectively.

## ASSOCIATED CONTENT

**Supporting Information**. The following files are available free of charge.

Supplemental materials and methods, supplementary tables and figures, supplemental data (PDF).

## AUTHOR INFORMATION

### Corresponding Author

James T. Stivers − Department of Pharmacology and Molecular Sciences, Johns Hopkins University School of Medicine 725 North Wolfe Street Baltimore, MD 21205, United States; orcid.org/0000-0003-2572- 7807; Phone: (410) 292-6844;

### Author

Benjamin Orris **−** Department of Pharmacology and Molecular Sciences, Johns Hopkins University School of Medicine 725 North Wolfe Street Baltimore, MD 21205, United States;

Shridhar Bhat **−** Department of Pharmacology and Molecular Sciences, Johns Hopkins University School of Medicine 725 North Wolfe Street Baltimore, MD 21205, United States;

Min Woo Sung − Medicine Design, Pfizer, Groton, CT 06340, USA;

Kevin W. Huynh – Medicine Design, Pfizer, Groton, CT 06340, USA;

Yingrong Xu – Medicine Design, Pfizer, Groton, CT 06340, USA;

Seungil Han − Medicine Design, Pfizer, Groton, CT 06340, USA;

Darren C. Johnson − Centers for Therapeutic Innovation (CTI), Pfizer, NY, NY 10016, USA

Benedikt Bosbach − Centers for Therapeutic Innovation (CTI), Pfizer, NY, NY 10016, USA

David J. Shields − Centers for Therapeutic Innovation (CTI), Pfizer, NY, NY 10016, USA

### Author Contributions

The manuscript was written through contributions of all authors. All authors have given approval to the final version of the manuscript.

## DATA AVAILIBILITY

All data pertaining to this study are archived at *to be filled in after acceptance for publication*. The final SAMHD1- ssRNA32 tetramer complex cryo-EM density maps and models are deposited in the Electron Microscopy Data Bank (EMDB) under accession code EMD-*to be filled in after acceptance for publication* and Protein Data Bank (PDB) under accession code *to be filled in after acceptance for publication*, respectively.

## ACKNOWLEDGMENTS

We acknowledge the helpful suggestions and insights provided by Matt Egleston.

