## Supplemental Information for "Guanine-containing ssDNA and RNA induce dimeric and tetrameric SAMHD1 in cryo-EM and binding studies"

### SUPPLEMENTAL METHODS

**Supplemental Method 1.** DynaFit Script used in Figure S2.

```
[task]
  task = fit
  data = equilibria

[mechanism]
  E.probe <===> E + probe : Kd.probe equil ; labeled ssNA equilibrium
  E.ssNA <===> E + ssNA : Kd.ssNA equil ; unlabeled ssNA equilibrium

[constants]
  Kd.probe = 2.13 ; 2.13 uM K_0.5
  Kd.ssNA = 2.13 ?; to be fitted

[concentrations]
  E = 1 ; 1 uM enzyme
  probe = 0.5 ; .5 uM labeled DNA

[responses]
  E.probe = 1 ?

[data]
  variable ssNA ; unlabeled competitor is being titrated
  file ./FigS2_comp/dT40_data.csv
  offset 0 ?

[output]
  directory ./FigS2_comp/dT40

[end]
```

**Supplemental Method S2.** Detailed protocol for proteomic analysis of SAMHD1 protein

Protein samples were combined with denaturation buffer (4% sodium deoxycholate, 100 mM Tris-HCl pH 8.5) to a total volume of 50 µL. Samples were shaken at 95 °C, 1000 rpm for 5 min. To each sample 5 µL reduction/alkylation buffer (100 mM Tris (2-carboxyethyl) phosphine, 400 mM chloroacetamide) was added, and samples were shaken at 45 °C, 1000 rpm for 5 min. SP3 (single-pot, solid-phase-enhanced sample-preparation) technology was used and digestion was performed on bead. A fresh 1:1 SP3 bead mix was prepared by combining equal volumes of stock beads (Sera-Mag SpeedBeads, GE Healthcare, cat. no 45152105050250 and cat. no 65152105050250) and washing the beads 3 times with ultrapure water. During the washes, beads were vortexed for 10 s and placed on the magnetic stand for 2 min to be collected. Supernatant was discarded. A total of 15 µL 50 mg/mL washed mixed beads were added to each sample before the addition of 70 µL ethanol for a final ethanol concentration of 50%. Samples were vortexed to mix, and were shaken at R.T., 1000 rpm for 10 min. The samples were spun briefly and placed on magnetic stand for 2 min. The beads were washed 3 times with 500 µL 80% ethanol. During the washes, beads were vortexed for 30 s then placed on magnetic stand for 2 min. After the washes, the beads were air dried for 30 s. The beads from each sample were resuspended in 50 µL 20 mM tetraethylammonium bromide (TEAB) pH 8.5 and were sonicated to mix. To each sample 5 µL trypsin (1 µg/µL in 20 mM TEAB) was added and samples were shaken at 37 °C, 1000 rpm overnight. The next day samples were sonicated and placed on magnetic stand for 2 min. Supernatant was transferred to a new tube and placed on magnetic stand again for 2 min. The supernatant was dried on a SpeedVac and reconstituted in 50 µL 1% formic acid, 2% acetonitrile for LC-MS/MS analysis.

LC-MS/MS data acquisition. Two µg of peptides were injected onto an EASY-Spray column (ES803A, 50 cm, 75 µm ID, 2 µm particle size, Thermo Fisher Scientific) heated to 60 °C. The peptides were separated using an EASY-nLC 1200 (Thermo Fisher Scientific) with a linear gradient from 10 to 32% B over 172 min at flow rate of 450 nL min$^{-1}$. Chromatography buffer A consisted of 0.1% formic acid in water, buffer B consisted of 0.1% formic acid in 80% acetonitrile. Mass spectra were acquired in positive ion mode using an Orbitrap Fusion Lumos Tribrid mass spectrometer (Thermo Fisher Scientific) equipped with an EASY-Spray NG ion source. $MS^1$ scan was conducted in the Orbitrap with a resolution of 120,000, maximum injection time of 50 ms, automatic gain control (AGC) target value of 4 x $10^5$ and mass range of 385-1415 Th. $MS^2$ was performed in the Orbitrap with a resolution of 50,000, using data-dependent acquisition mode with 3 s cycle time, higher-energy collisional dissociation (HCD, CE=30%), normal m/z scan range mode, maximum injection time of 86 ms, AGC target of 2 x $10^4$, dynamic exclusion (DE=90 s), and isolation window of 0.5 Th by quadrupole isolation. The $MS^1$ scans were in profile mode and the $MS^2$ scans were in centroid mode.

SAMHD1 immunoprecipitation. Two T-175 flasks were seeded with 80x$10^6$ THP1 Dual cells (InvivoGen, cat. no. thpd-nfis) in 50 mL of complete medium (RPMI 1640 + 10% heat inactivated fetal bovine serum + 100 U/mL Penicillin/Streptomycin). 4 mM Hydroxyurea was added to each flask and flasks were incubated for 2 h in a humidified incubator at 37 °C and 5% CO2 to induce replication stress. Cells from the treated flasks were pooled and nuclear fractions were extracted using a Subcellular Fractionation Kit for Tissues (Thermo Fisher Scientific cat. no. 87790) according to the manufacturer's instructions. The Nuclear extract was normalized to 1 mg/mL and pre-cleared by incubating with Protein A Magnetic Beads (Cell Signaling, cat. no. 73778) according to the manufacturer's instructions. IgG XP® Isotype Control (Cell Signaling, cat. on. 3900), phospho-SAMHD1 (Cell Signaling Technology, cat. no. 89930), SAMHD1 N-terminal (Abcam, cat. no. Ab191420), and SAMHD1 (Bethyl Laboratories, cat. no. 303-690A) antibodies were added to the nuclear extracts (6 μg antibody/mg lysate) and incubated overnight at 4 °C with rotation. Protein A Magnetic Beads were added to the immunocomplexes and immunoprecipitated according to manufacturer's instructions. Washed bead pellets were resuspended in ice cold HEPES buffer (20 mM, pH ~8.5) and 1/4 volume bead pellets were aliquoted for western blot analysis. Western blot samples were prepared by resuspending bead pellets in 30 μL 3X Blue Loading Buffer (Cell Signaling Technology, cat. no. 56036S) supplemented with DTT (125 mM final concentration). The samples were then boiled at 100 °C for 15 min and subsequently analyzed by western blot. LC-MS samples were prepared by adding 50 μL 6 M Urea supplemented with 10 mM DTT (final concentration) to the remaining bead pellets and incubating the samples for 20 min at 37 °C with shaking. Iodoacetamide (20 mM, final concentration) was added, and the samples were incubated for 30 min at room temperature with shaking, protected from light. After 30 min, the protein was digested with 2 ug Trypsin/Lys-C Protease Mix (Thermo Fisher Scientific, cat. no. A40007) for 9 h at room temperature. Samples were then analyzed by LC-MS.

LC-MS/MS data analysis. Raw data files were searched using Proteome Discoverer 3.0 (Thermo Fisher Scientific) with Sequest HT against the *Homo sapiens* database (SwissProt TaxID=9606_and_subtaxonomies), the Escherichia coli database (sp_canonical TaxID=562_and_subtaxonomies) and an in-house contaminants database. The searches were performed with the following parameters: MS tolerance 10 ppm, MS/MS tolerance 0.02 Da, trypsin digestion with 2 missed cleavages allowed, fixed carbamidomethyl modification of cysteine, variable methionine oxidation, variable asparagine deamidation, variable acetylation of protein N-terminus, variable methionine loss of protein N-terminus and variable acetylation in addition to methionine loss of protein N-terminus. Only peptides with high confidence (false discovery rate or FDR < 1%) and proteins with medium or high confidence (FDR < 5%) were used. Only proteins with at least 2 unique peptides were used in the analysis.

### SUPPLEMENTAL TABLES

**Supplemental Table S1.** Sequences of nucleic acid constructs.

| Name | RNA or DNA | Sequence |
|---|---|---|
| 5′ Guanine Specificity | | |
| 5′FAM-dT$_{40}$ | DNA | 5’-/FAM/TTT TTT TTT TTT TTT TTT TTT TTT TTT TTT TTT TTT TTT T-3’ |
| 5′FAM-dT**dA**dT$_{38}$ | DNA | 5’-/FAM/TAT TTT TTT TTT TTT TTT TTT TTT TTT TTT TTT TTT TTT T-3’ |
| 5′FAM-dT**dX**dT$_{38}$ | DNA | 5’-/FAM/TXT TTT TTT TTT TTT TTT TTT TTT TTT TTT TTT TTT TTT T-3’ |
| 5′FAM-dT**dG**dT$_{38}$ | DNA | 5’-/FAM/TGT TTT TTT TTT TTT TTT TTT TTT TTT TTT TTT TTT TTT T-3’ |
| dT$_{38}$**dX**dT-3’FAM | DNA | 5’-TTT TTT TTT TTT TTT TTT TTT TTT TTT TTT TTT TTT TTX T/FAM/-3’ |
| dT$_{38}$**dG**dT-3’FAM | DNA | 5’-TTT TTT TTT TTT TTT TTT TTT TTT TTT TTT TTT TTT TTG T/FAM/-3’ |
| dT-6FAMdT-dT$_{38}$ | DNA | 5’-T/6FAMdT/TT TTT TTT TTT TTT TTT TTT TTT TTT TTT TTT TTT TTT T-3’ |
| **dG**-6FAMdT-dT$_{38}$ | DNA | 5’-G/6FAMdT/TT TTT TTT TTT TTT TTT TTT TTT TTT TTT TTT TTT TTT T-3’ |
| 5′FAM-U$_{40}$ | RNA | 5’-/FAM/UUU UUU UUU UUU UUU UUU UUU UUU UUU UUU UUU UUU UUU U-3’ |
| 5′FAM-U**A**U$_{38}$ | RNA | 5’-/FAM/UAU UUU UUU UUU UUU UUU UUU UUU UUU UUU UUU UUU UUU U-3’ |
| 5′FAM-U**G**U$_{38}$ | RNA | 5’-/FAM/UGU UUU UUU UUU UUU UUU UUU UUU UUU UUU UUU UUU UUU U-3’ |
| U$_{38}$**G**U-3’FAM | RNA | 5’-UUU UUU UUU UUU UUU UUU UUU UUU UUU UUU UUU UUU UUG U/FAM/-3’ |
| Positional Effects | | |
| 5′FAM-**dG**dT$_{39}$ | DNA | 5’-/FAM/GTT TTT TTT TTT TTT TTT TTT TTT TTT TTT TTT TTT TTT T-3’ |
| 5′FAM-dT$_{2}$**dG**dT$_{37}$ | DNA | 5’-/FAM/TTG TTT TTT TTT TTT TTT TTT TTT TTT TTT TTT TTT TTT T-3’ |
| 5′FAM-dT$_{3}$**dG**dT$_{36}$ | DNA | 5’-/FAM/TTT GTT TTT TTT TTT TTT TTT TTT TTT TTT TTT TTT TTT T-3’ |
| 5′FAM-dT$_{4}$**dG**dT$_{35}$ | DNA | 5’-/FAM/TTT TGT TTT TTT TTT TTT TTT TTT TTT TTT TTT TTT TTT T-3’ |
| 5′FAM-dT$_{6}$**dG**dT$_{33}$ | DNA | 5’-/FAM/TTT TTT GTT TTT TTT TTT TTT TTT TTT TTT TTT TTT TTT T-3’ |
| 5′FAM-dT$_{9}$**dG**dT$_{30}$ | DNA | 5’-/FAM/TTT TTT TTT GTT TTT TTT TTT TTT TTT TTT TTT TTT TTT T-3’ |
| 5′FAM-dT$_{12}$**dG**dT$_{27}$ | DNA | 5’-/FAM/TTT TTT TTT TTT GTT TTT TTT TTT TTT TTT TTT TTT TTT T-3’ |
| 5′FAM-dT$_{25}$**dG**dT$_{14}$ | DNA | 5’-/FAM/TTT TTT TTT TTT TTT TTT TTT TTT TGT TTT TTT TTT TTT T-3’ |
| 5′FAM-dT$_{38}$**dG**dT | DNA | 5’-/FAM/TTT TTT TTT TTT TTT TTT TTT TTT TTT TTT TTT TTT TTG T-3’ |
| 5′FAM-U$_{12}$**G**U$_{27}$ | RNA | 5’-/FAM/UUU UUU UUU UUU GUU UUU UUU UUU UUU UUU UUU UUU UUU U-3’ |
| 5′FAM-U$_{25}$**G**U$_{14}$ | RNA | 5’-/FAM/UUU UUU UUU UUU UUU UUU UUU UUU UGU UUU UUU UUU UUU U-3’ |
| Length Effects | | |
| 5′FAM-**dG**dT$_{29}$ | DNA | 5’-/FAM/GTT TTT TTT TTT TTT TTT TTT TTT TTT TTT-3’ |
| 5′FAM-**dG**dT$_{24}$ | DNA | 5’-/FAM/GTT TTT TTT TTT TTT TTT TTT TTT T-3’ |
| 5′FAM-**dG**dT$_{19}$ | DNA | 5’-/FAM/GTT TTT TTT TTT TTT TTT TT-3’ |
| 5′FAM-**dG**dT$_{14}$ | DNA | 5’-/FAM/GTT TTT TTT TTT TTT-3’ |
| 5′FAM-**dG**dT$_{9}$ | DNA | 5’-/FAM/GTT TTT TTT T-3’ |
| Mixed Sequence 32mers | | |
| 5′FAM-ssDNA$_{32}$ | DNA | 5’ - /FAM/ CAC TAT CGG AAA AAG GGC GAC ACG GAT ATG TT – 3’ |
| 5′FAM-ssRNA$_{32}$ | RNA | 5’ - /FAM/ CAC UAU CGG AAA AAG GGC GAC ACG GAU AUG UU – 3’ |
| Two-Guanine Spacing | | |
| 5′FAM-**dG**dT$_{3}$**dG**dT$_{35}$ | DNA | 5’-/FAM/GTT TGT TTT TTT TTT TTT TTT TTT TTT TTT TTT TTT TTT T-3’ |
| 5′FAM-**dG**dT$_{8}$**dG**dT$_{30}$ | DNA | 5’-/FAM/GTT TTT TTT GTT TTT TTT TTT TTT TTT TTT TTT TTT TTT T-3’ |
| 5′FAM-**dG**dT$_{13}$**dG**dT$_{25}$ | DNA | 5’-/FAM/GTT TTT TTT TTT TTG TTT TTT TTT TTT TTT TTT TTT TTT T-3’ |
| 5′FAM-**dG**dT$_{18}$**dG**dT$_{20}$ | DNA | 5’-/FAM/GTT TTT TTT TTT TTT TTT TGT TTT TTT TTT TTT TTT TTT T-3’ |
| 5′FAM-**dG**dT$_{23}$**dG**dT$_{15}$ | DNA | 5’-/FAM/GTT TTT TTT TTT TTT TTT TTT TTT GTT TTT TTT TTT TTT T-3’ |

| | | |
|---|---|---|
| 5′FAM-**dG**dT$_{28}$**dG**dT$_{10}$ | DNA | 5’-/FAM/GTT TTT TTT TTT TTT TTT TTT TTT TTT TTG TTT TTT TTT T-3’ |
| 5′FAM-**dG**dT$_{33}$**dG**dT$_{5}$ | DNA | 5’-/FAM/GTT TTT TTT TTT TTT TTT TTT TTT TTT TTT TTT TGT TTT T-3’ |
| 5′FAM-**dG**dT$_{38}$**dG** | DNA | 5’-/FAM/GTT TTT TTT TTT TTT TTT TTT TTT TTT TTT TTT TTT TTT G-3’ |
| 5′FAM-**G**U$_{3}$**G**U$_{35}$ | RNA | 5’-/FAM/GUU UGU UUU UUU UUU UUU UUU UUU UUU UUU UUU UUU UUU U-3’ |
| 5′FAM-**G**U$_{8}$**G**U$_{30}$ | RNA | 5’-/FAM/GUU UUU UUU GUU UUU UUU UUU UUU UUU UUU UUU UUU UUU U-3’ |
| 5′FAM-**G**U$_{13}$**G**U$_{25}$ | RNA | 5’-/FAM/GUU UUU UUU UUU UUG UUU UUU UUU UUU UUU UUU UUU UUU U-3’ |
| 5′FAM-**G**U$_{18}$**G**U$_{20}$ | RNA | 5’-/FAM/GUU UUU UUU UUU UUU UUU UGU UUU UUU UUU UUU UUU UUU U-3’ |
| 5′FAM-**G**U$_{23}$**G**U$_{15}$ | RNA | 5’-/FAM/GUU UUU UUU UUU UUU UUU UUU UUU GUU UUU UUU UUU UUU U-3’ |
| 5′FAM-**G**U$_{28}$**G**U$_{10}$ | RNA | 5’-/FAM/GUU UUU UUU UUU UUU UUU UUU UUU UUU UUG UUU UUU UUU U-3’ |
| 5′FAM-**G**U$_{33}$**G**U$_{5}$ | RNA | 5’-/FAM/GUU UUU UUU UUU UUU UUU UUU UUU UUU UUU UUU UGU UUU U-3’ |
| 5′FAM-**G**U$_{38}$**G** | RNA | 5’-/FAM/GUU UUU UUU UUU UUU UUU UUU UUU UUU UUU UUU UUU UUU G-3’ |

**Supplemental Table S2.** Binding parameters of nucleic acid constructs[a]

| Name | RNA or DNA | $K_{0.5}$ (µM) | $A_{max}$ | *n* |
|---|---|---|---|---|
| 5′ Guanine Specificity | | | | |
| 5′FAM-dT$_{40}$ | DNA | 2.13 ± 0.38 | 0.06 ± 0.006 | 2.18 ± 0.67 |
| 5′FAM-dT**dA**dT$_{38}$ | DNA | 1.56 ± 0.19 | 0.06 ± 0.003 | 1.88 ± 0.35 |
| 5′FAM-dT**dX**dT$_{38}$ | DNA | 0.68 ± 0.06 | 0.19 ± 0.005 | 1.86 ± 0.26 |
| 5′FAM-dT**dG**dT$_{38}$ | DNA | 0.35 ± 0.02 | 0.24 ± 0.003 | 2.49 ± 0.24 |
| dT$_{38}$**dX**dT-3′FAM | DNA | 1.74 ± 0.64 | 0.06 ± 0.011 | 1.29 ± 0.43 |
| dT$_{38}$**dG**dT-3′FAM | DNA | 1.59 ± 0.13 | 0.10 ± 0.004 | 1.85 ± 0.23 |
| dT-6FAMdT-dT$_{38}$ | DNA | 1.20 ± 0.12 | 0.10 ± 0.004 | 2.21 ± 0.41 |
| **dG**-6FAMdT-dT$_{38}$ | DNA | 0.26 ± 0.01 | 0.25 ± 0.002 | 2.49 ± 0.20 |
| 5′FAM-U$_{40}$ | RNA | 1.29 ± 0.13 | 0.07 ± 0.004 | 1.96 ± 0.35 |
| 5′FAM-U**A**U$_{38}$ | RNA | 1.64 ± 0.19 | 0.08 ± 0.004 | 1.63 ± 0.24 |
| 5′FAM-U**G**U$_{38}$ | RNA | 0.16 ± 0.01 | 0.24 ± 0.002 | 2.06 ± 0.13 |
| U$_{38}$**G**U-3′FAM | RNA | 1.10 ± 0.14 | 0.13 ± 0.007 | 1.35 ± 0.19 |
| Positional Effects | | | | |
| 5′FAM-**dG**dT$_{39}$ | DNA | 0.25 ± 0.01 | 0.24 ± 0.002 | 2.54 ± 0.20 |
| 5′FAM-dT$_{2}$**dG**dT$_{37}$ | DNA | 0.33 ± 0.02 | 0.25 ± 0.002 | 2.45 ± 0.14 |
| 5′FAM-dT$_{3}$**dG**dT$_{36}$ | DNA | 0.32 ± 0.01 | 0.24 ± 0.002 | 2.37 ± 0.12 |
| 5′FAM-dT$_{4}$**dG**dT$_{35}$ | DNA | 0.37 ± 0.01 | 0.23 ± 0.002 | 2.33 ± 0.13 |
| 5′FAM-dT$_{6}$**dG**dT$_{33}$ | DNA | 0.39 ± 0.03 | 0.16 ± 0.003 | 1.84 ± 0.19 |
| 5′FAM-dT$_{9}$**dG**dT$_{30}$ | DNA | 0.52 ± 0.05 | 0.11 ± 0.003 | 2.02 ± 0.32 |
| 5′FAM-dT$_{12}$**dG**dT$_{27}$ | DNA | 0.64 ± 0.06 | 0.07 ± 0.003 | 1.97 ± 0.35 |
| 5′FAM-dT$_{25}$**dG**dT$_{14}$ | DNA | 1.17 ± 0.13 | 0.07 ± 0.003 | 2.05 ± 0.39 |
| 5′FAM-dT$_{38}$**dG**dT | DNA | 0.97 ± 0.08 | 0.13 ± 0.004 | 2.17 ± 0.32 |
| 5′FAM-U$_{12}$**G**U$_{27}$ | RNA | 0.41 ± 0.03 | 0.08 ± 0.002 | 1.48 ± 0.14 |
| 5′FAM-U$_{25}$**G**U$_{14}$ | RNA | 0.77 ± 0.07 | 0.08 ± 0.003 | 1.58 ± 0.20 |
| Length Effects | | | | |
| 5′FAM-**dG**dT$_{29}$ | DNA | 0.34 ± 0.02 | 0.22 ± 0.004 | 2.56 ± 0.34 |
| 5′FAM-**dG**dT$_{24}$ | DNA | 0.45 ± 0.03 | 0.23 ± 0.004 | 2.12 ± 0.25 |
| 5′FAM-**dG**dT$_{19}$ | DNA | 0.58 ± 0.05 | 0.22 ± 0.007 | 2.26 ± 0.42 |
| 5′FAM-**dG**dT$_{14}$ | DNA | 0.91 ± 0.08 | 0.21± 0.007 | 2.07 ± 0.32 |
| 5′FAM-**dG**dT$_{9}$ | DNA | 1.87 ± 0.24 | 0.18 ± 0.011 | 1.94 ± 0.38 |
| Mixed Sequence 32mers | | | | |
| 5′FAM-ssDNA$_{32}$ | DNA | 0.27 ± 0.01 | 0.17 ± 0.002 | 1.97 ± 0.15 |
| 5′FAM-ssRNA$_{32}$ | RNA | 0.15 ± 0.01 | 0.20 ± 0.003 | 3.10 ± 0.42 |
| Two-Guanine Spacing | | | | |
| 5′FAM-**dG**dT$_{3}$**dG**dT$_{35}$ | DNA | 0.17 ± 0.007 | 0.21 ± 0.002 | 2.86 ± 0.26 |
| 5′FAM-**dG**dT$_{8}$**dG**dT$_{30}$ | DNA | 0.18 ± 0.005 | 0.21 ± 0.002 | 2.59 ± 0.15 |
| 5′FAM-**dG**dT$_{13}$**dG**dT$_{25}$ | DNA | 0.19 ± 0.008 | 0.23 ± 0.003 | 2.42 ± 0.21 |
| 5′FAM-**dG**dT$_{18}$**dG**dT$_{20}$ | DNA | 0.16 ± 0.006 | 0.23 ± 0.002 | 2.40 ± 0.18 |
| 5′FAM-**dG**dT$_{23}$**dG**dT$_{15}$ | DNA | 0.15 ± 0.008 | 0.23 ± 0.003 | 2.62 ± 0.33 |
| 5′FAM-**dG**dT$_{28}$**dG**dT$_{10}$ | DNA | 0.15 ± 0.008 | 0.23 ± 0.003 | 2.42 ± 0.26 |
| 5′FAM-**dG**dT$_{33}$**dG**dT$_{5}$ | DNA | 0.16 ± 0.008 | 0.22 ± 0.002 | 2.46 ± 0.19 |
| 5′FAM-**dG**dT$_{38}$**dG** | DNA | 0.12 ± 0.008 | 0.22 ± 0.003 | 1.94 ± 0.18 |
| 5′FAM-**G**U$_{3}$**G**U$_{35}$ | RNA | 0.09 ± 0.004 | 0.24 ± 0.002 | 2.20 ± 0.18 |
| 5′FAM-**G**U$_{8}$**G**U$_{30}$ | RNA | 0.11 ± 0.004 | 0.25 ± 0.002 | 2.19 ± 0.12 |
| 5′FAM-**G**U$_{13}$**G**U$_{25}$ | RNA | 0.12 ± 0.005 | 0.25 ± 0.003 | 2.40 ± 0.21 |

| | | | | |
|---|---|---|---|---|
| 5′FAM-**G**U$_{18}$**G**U$_{20}$ | RNA | 0.12 ± 0.005 | 0.26 ± 0.003 | 2.49 ± 0.23 |
| 5′FAM-**G**U$_{23}$**G**U$_{15}$ | RNA | 0.10 ± 0.004 | 0.26 ± 0.003 | 2.34 ± 0.20 |
| 5′FAM-**G**U$_{28}$**G**U$_{10}$ | RNA | 0.11 ± 0.005 | 0.26 ± 0.003 | 2.01 ± 0.16 |
| 5′FAM-**G**U$_{33}$**G**U$_{5}$ | RNA | 0.10 ± 0.005 | 0.26 ± 0.003 | 2.08 ± 0.19 |
| 5′FAM-**G**U$_{38}$**G** | RNA | 0.11 ± 0.004 | 0.26 ± 0.003 | 2.45 ± 0.20 |

[a]$K_{0.5}$ is obtained from the Hill equation (eq 2), $n$ is the Hill coefficient and $A_{max}$ is the maximal observed fluorescence anisotropy.

**Supplemental Table S3.** Cryo-EM data collection, 3D reconstruction and model building statistics.

| | | |
|---|---|---|
| **Data Collection** | | |
| **Microscope** | Titan Krios | |
| **Camera** | Falcon4i | |
| **Magnification** | 165 000 | |
| **Voltage (kV)** | 300 | |
| **Total dose ($e^-$/Å$^2$)** | 40 | |
| **Defocus range (µm)** | -0.6 to -2.8 | |
| **Number of fractions** | 40 | |
| **Movies** | 26702 | |
| **Pixel spacing (Å)** | 0.733 | |
| **Symmetry imposed** | *C*1 | |
| **FSC threshold** | 0.143 | |
| | **T*-closed** | **T*-open** |
| **Map pixel size (Å/pxl)** | 1.20 | 1.20 |
| **Real-space correlation (mask)** | 0.61 | 0.5 |
| **MolProbity score** | 1.79 | 1.75 |
| **Clash score** | 6.0 | 6.70 |
| **Cβ outliers (%)** | 0 | 0 |
| **Rotamer (%)** | 0.53 | 0 |
| **Ramachandran plot (%)** | | |
| **Favored** | 92.64 | 94.36 |
| **Allowed** | 7.36 | 5.64 |
| **Outliers** | 0 | 0 |
| **RMS deviations** | | |
| **Bond length** | 0.002 | 0.005 |
| **Bond angles** | 0.525 | 0.563 |

**Supplemental Table S4.** Mass spectrometry analysis of trace proteins present in SAMHD1 protein preparations[a]

| Purification step | Total proteins | Human | E. coli | RNase | DNase |
|---|---|---|---|---|---|
| Post-Ni-NTA | 424 | 21 | 403 | 3 (RpsP, PNP, RnE) | 1 (UvrA) |
| Post-SP sepharose | 211 | 13 | 198 | 2 (RpsP, RnE) | 1 (UvrA) |
| Post-SEC | 78 | 6 | 72 | 1 (RnE) | 1 (UvrA) |

[a]SAMHD1 with a 10x N-terminal histidine tag was purified as described in methods.

### SUPPLEMENTAL FIGURES

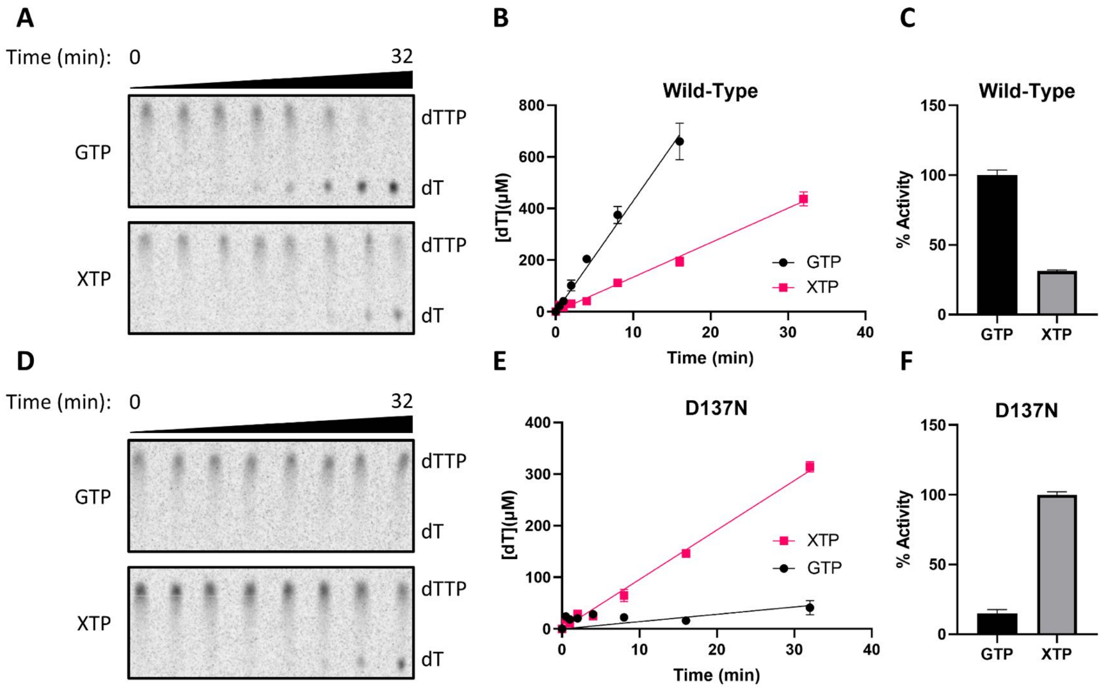

**Supplemental Figure S1. SAMHD1 D137N is activated by XTP instead of GTP. (A)** Reversed-phase TLC chromatograms of the hydrolysis of dTTP (1 mM) by wild-type SAMHD1 (0.5 µM) in the presence of GTP (0.5 mM) (top) or XTP (0.5 mM) (bottom) over the course of 32 minutes. **(B)** Linear regression analysis of [dT] vs. time in the initial, linear phase of the reactions in (A). Data from the GTP-containing reaction is plotted as black circles and data from the XTP-containing reaction is plotted as pink rectangles. **(C)** Normalized reaction rates from (B), where 0 % activity is 0 µM dT/minute, and 100 % activity is the reaction rate attained with GTP activator. **(D)** Reversed-phase TLC chromatograms of the hydrolysis of dTTP (1 mM) by SAMHD1 D137N (0.5 µM) in the presence of GTP (0.5 mM) (top) or XTP (0.5 mM) (bottom) over the course of 32 minutes. **(E)** Linear regression analysis of [dT] vs. time of the reactions in (D). Data from the GTP-containing reaction is plotted as black circles and data from the XTP-containing reaction is plotted as pink rectangles. **(F)** Normalized reaction rates from (E), where 0 % activity is 0 µM dT/minute, and 100 % activity is the reaction rate attained with XTP activator.

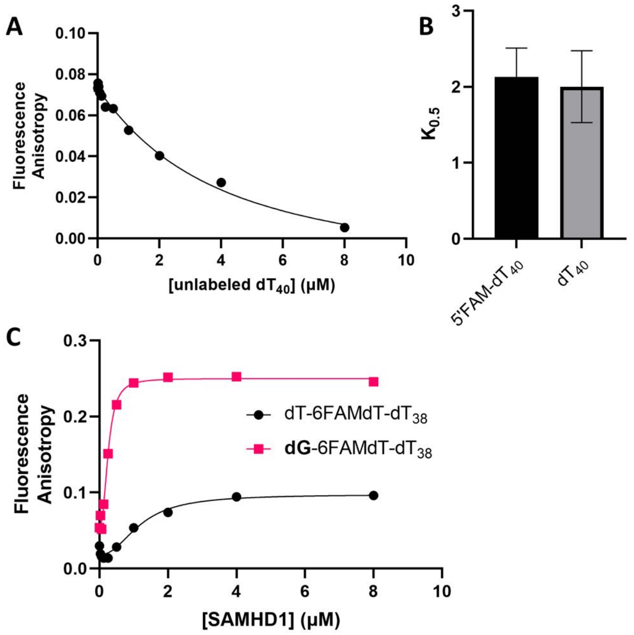

**Supplemental Figure S2. 5'-FAM labels do not affect ssDNA binding affinity and are not responsible for 5' G specificity.** (**A**) Displacement of 5'FAM-$dT_{40}$ (0.5 µM) from SAMHD1 (1 µM) by increasing concentrations of unlabeled $dT_{40}$. Error bars indicate standard error of three independent replicate measurements at each unlabeled $dT_{40}$ concentration. Data was subject to numerical regression fit using DynaFit (**Supplemental Method S1**). (**B**) Comparison of $K_{0.5}$ between 5'FAM-$dT_{40}$ (as determined by least-squares regression fit to equation 2, see **Fig. 1B**) and unlabeled $dT_{40}$ (as determined by Dynafit, see A). (**C**) Binding of SAMHD1 to dT-6FAMdT-$dT_{38}$ (black circles) and to **dG**-6FAMdT-$dT_{38}$ (pink squares). Error bars indicate standard error of three independent replicate measurements at each SAMHD1 concentration.

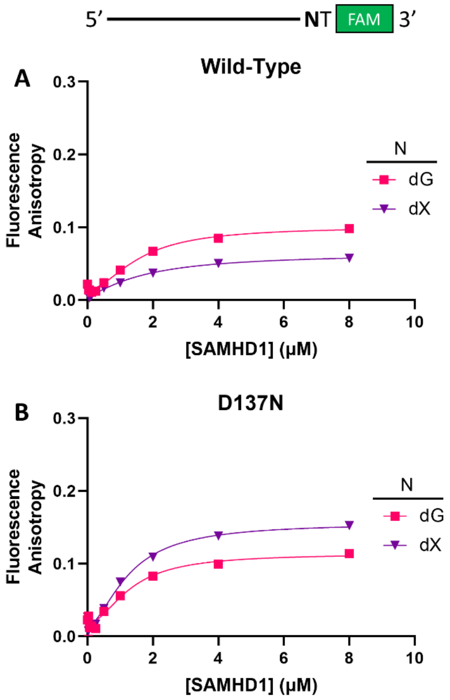

**Supplemental Figure S3. G or X positioned adjacent to the 3′ end of an oligonucleotide results in weak A1 site binding.** (**A**) Binding of wild-type SAMHD1 to two $dT_{38}$**N**dT-3′FAM (50 nM) where N = dG (pink squares) or N = dX (purple triangles). Error bars indicate standard error of three independent replicate measurements at each SAMHD1 concentration. (**B**) Binding of SAMHD1 D137N to the same two oligonucleotides as in (A). Error bars indicate standard error of three independent replicate measurements at each SAMHD1 concentration.

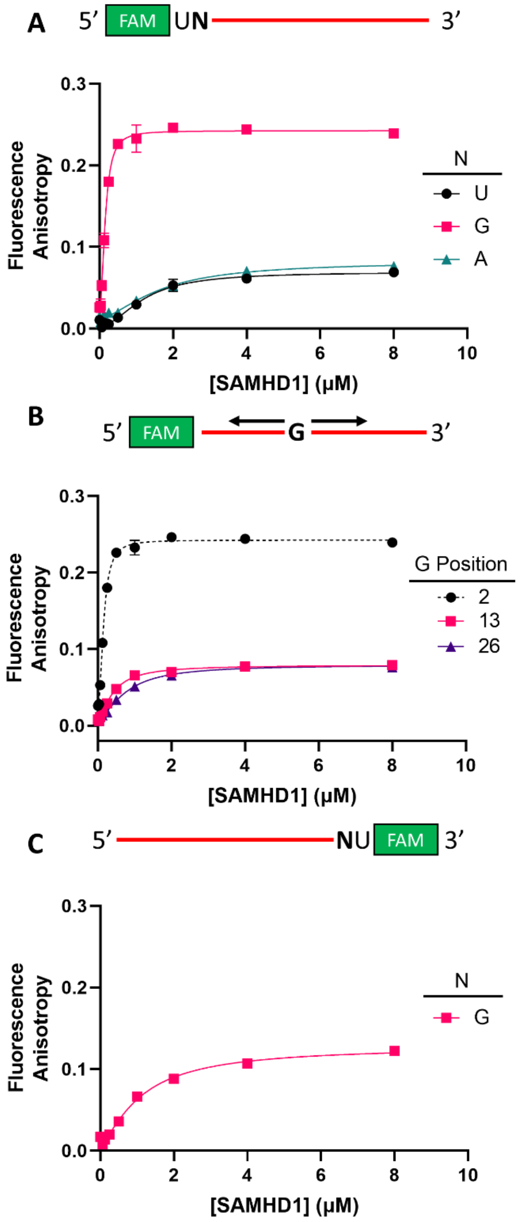

**Supplemental Figure S4. Specificity for guanine bases in ssRNA.** (**A**) Binding of SAMHD1 to 5′FAM-U**N**$U_{38}$ (50 nM) where N = U (black), G (pink), or A (green). Error bars indicate standard error of three independent replicate measurements at each SAMHD1 concentration. (**B**) Binding of SAMHD1 to 5′FAM-$U_{12}$**G**$U_{27}$ (pink) and 5′FAM-$U_{25}$**G**$U_{14}$ (purple) (50 nM each) in comparison with to 5′FAM-U**N**$U_{38}$ (black, from A). Error bars indicate standard error of three independent replicate measurements at each SAMHD1 concentration. (**C**) Binding of wild-type SAMHD1 to a $U_{38}$**G**U-3′FAM. Error bars indicate standard error of three independent replicate measurements at each SAMHD1 concentration.

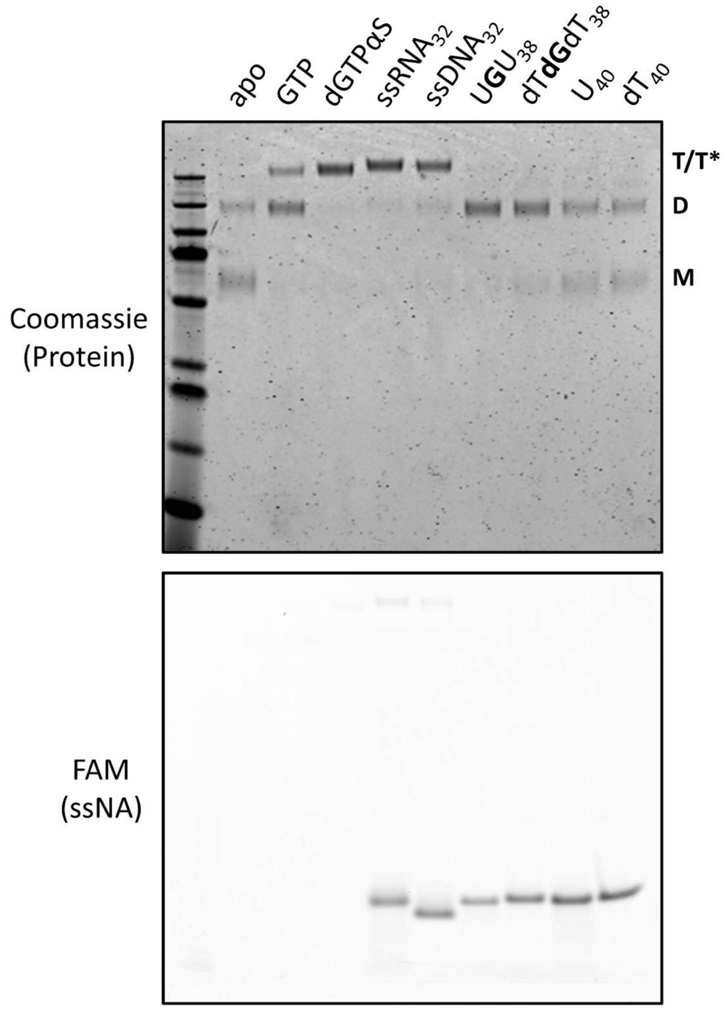

**Supplemental Figure S5. Nucleic acids are not efficiently crosslinked to SAMHD1 during glutaraldehyde crosslinking.** SAMHD1 (1 µM) was incubated alone, or in the presence of GTP (50 µM), dGTPαS (100 µM), or in the presence of the delineated 5'FAM-labeled ssRNA and ssDNA oligonucleotides (1 µM) for 10 minutes, then crosslinked for 10 minutes with glutaraldehyde (50 mM). Oligonucleotides were visualized by imaging the PAGE gel using FAM-fluorescence detection (bottom). SAMHD1 was visualized by staining the gel with Coomassie R250 dye and then imaged with fluorescent Coomassie detection (top). The small amount of $ssRNA_{32}$ and $ssDNA_{32}$ in the lower image that comigrate with the enzyme tetramer is less than 5% of the total ssNA present.

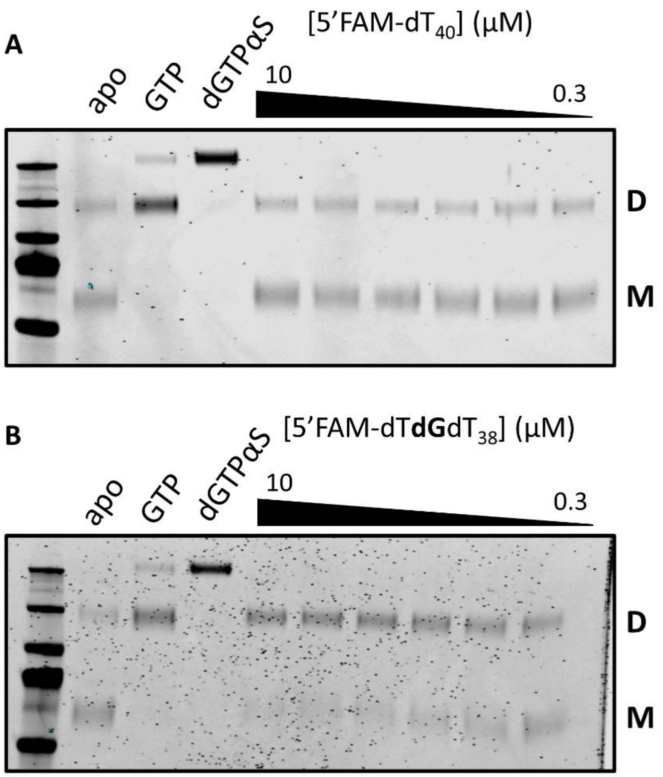

**Supplemental Figure S6. Guanine bases in ssDNA induce SAMHD1 dimerization in a concentration-dependent manner.** GAXL crosslinking experiments were carried out as in Fig. 3. (**A**) Oligomeric states of SAMHD1 induced by various concentrations of 5′FAM-$dT_{40}$. Lanes from left to right reflect serial two-fold dilutions in the range 10 to 0.3 μM. (**B**) Oligomeric states of SAMHD1 induced by various concentrations of 5′FAM-dT**dG**$dT_{38}$. Lanes from left to right reflect serial two-fold dilutions in the range 10 to 0.3 μM.

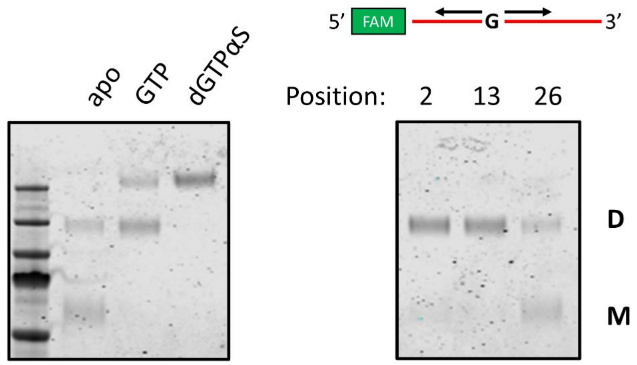

**Supplemental Figure S7. Guanine base positional dependence of SAMHD1 dimerization in the context of ssRNA.** GAXL crosslinking experiments were carried out as described in Fig. 3. Oligomeric states induced by 5'FAM-labeled U homopolymer ssRNA 40mers (1 μM) as a function of the position of the dG base within the sequence.

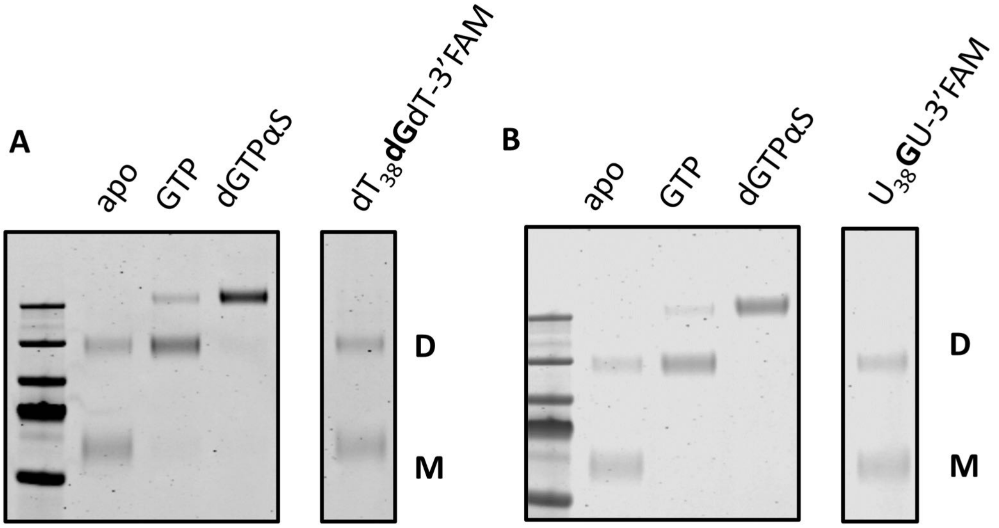

**Supplemental Figure S8. SAMHD1 dimerization is not induced by 3' end-proximate guanine bases in ssDNA and ssRNA.** GAXL crosslinking experiments were carried out as in Fig. 3. (**A**) Oligomeric states induced by $dT_{38}$**dG**dT-3'FAM (1 μM). (**B**) Oligomeric states induced by $U_{38}$**G**U-3'FAM (1 μM).

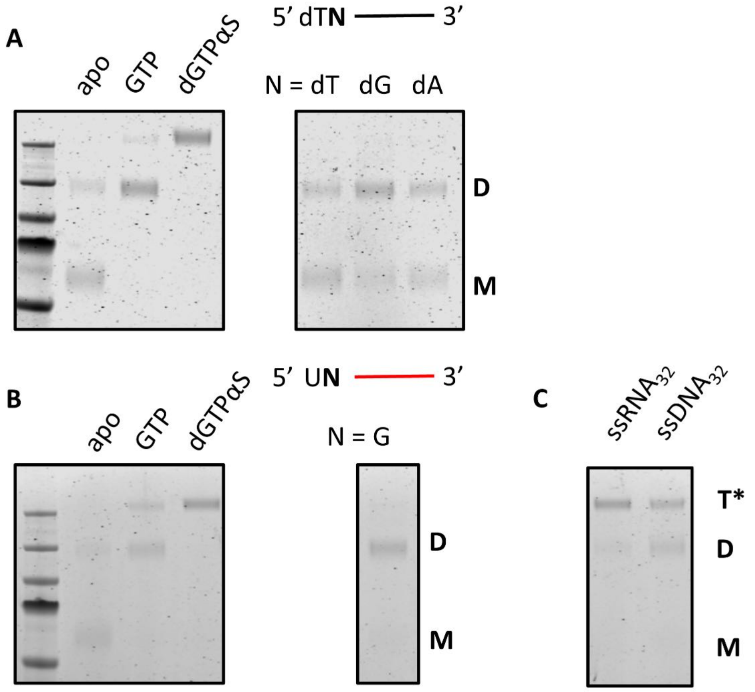

**Supplemental Figure S9. Guanine-dependent dimerization and tetramerization occur in the absence of a 5′-FAM label**. GAXL crosslinking experiments were carried out as in Fig. 3. (**A**) Oligomeric states of SAMHD1 induced by 1 µM dT**dN**dT$_{38}$ (where N = dT, dG or dA). (**B**) Oligomeric states of SAMHD1 induced by 1 µM U**G**U$_{38}$. (**C**) Oligomeric states of SAMHD1 induced by 1 µM unlabeled ssRNA$_{32}$ and ssDNA$_{32}$.

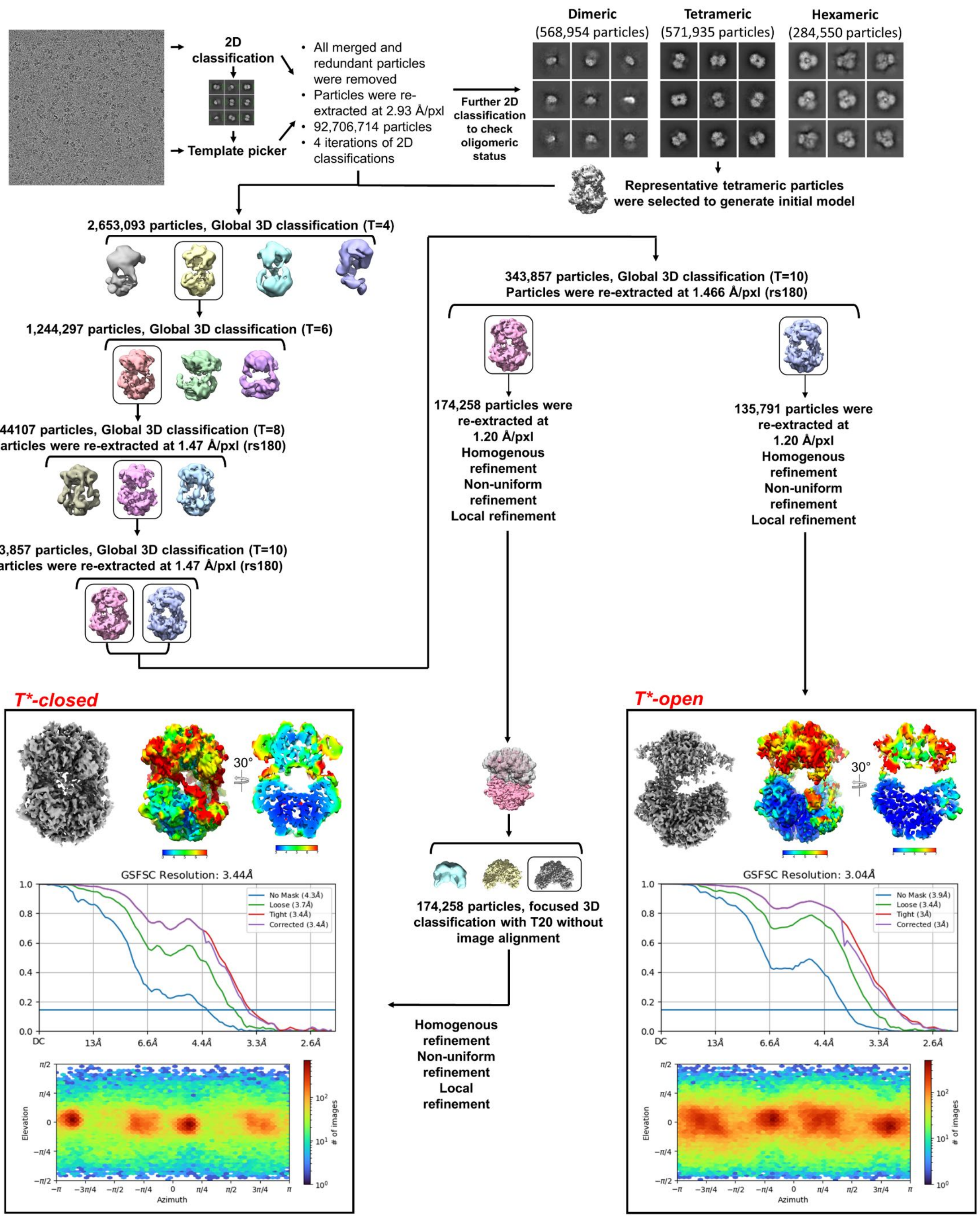

**Supplemental Figure S10. cryoEM Data processing workflow.** 2D and 3D structural classification steps are outlined and the further 3D refinements of the T*closed and T*open conformations are shown. The Fourier Shell Correlation (FSC) curves between two independently refined half maps for T*closed and T*open are shown in the bottom left and right panels, respectively. Local resolution for T*closed and T*open (overall and dissected) and particle distributions are indicated.

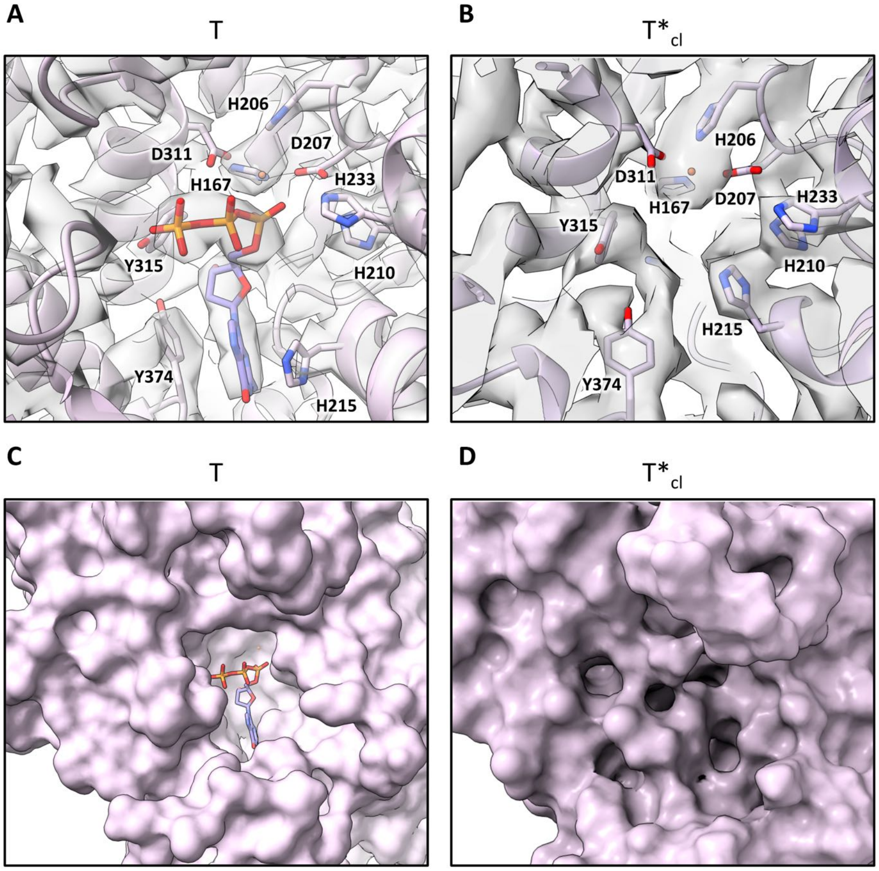

**Supplemental Figure S11. Comparison of T and T*$_{cl}$ active sites.** (**A**) Active site of the canonical SAMHD1 tetramer as induced by dGTPαS (PDB: 7UJN). An iron atom and a dGTPαS are modeled into the electron density. (**B**) Active site of T*$_{cl}$, contains electron density for the iron, but is devoid of any nucleotide density. (**C**) Surface model of the active site cleft of the canonical SAMHD1 tetramer. (**D**) Surface model of the occluded active site cleft of T*$_{cl}$.

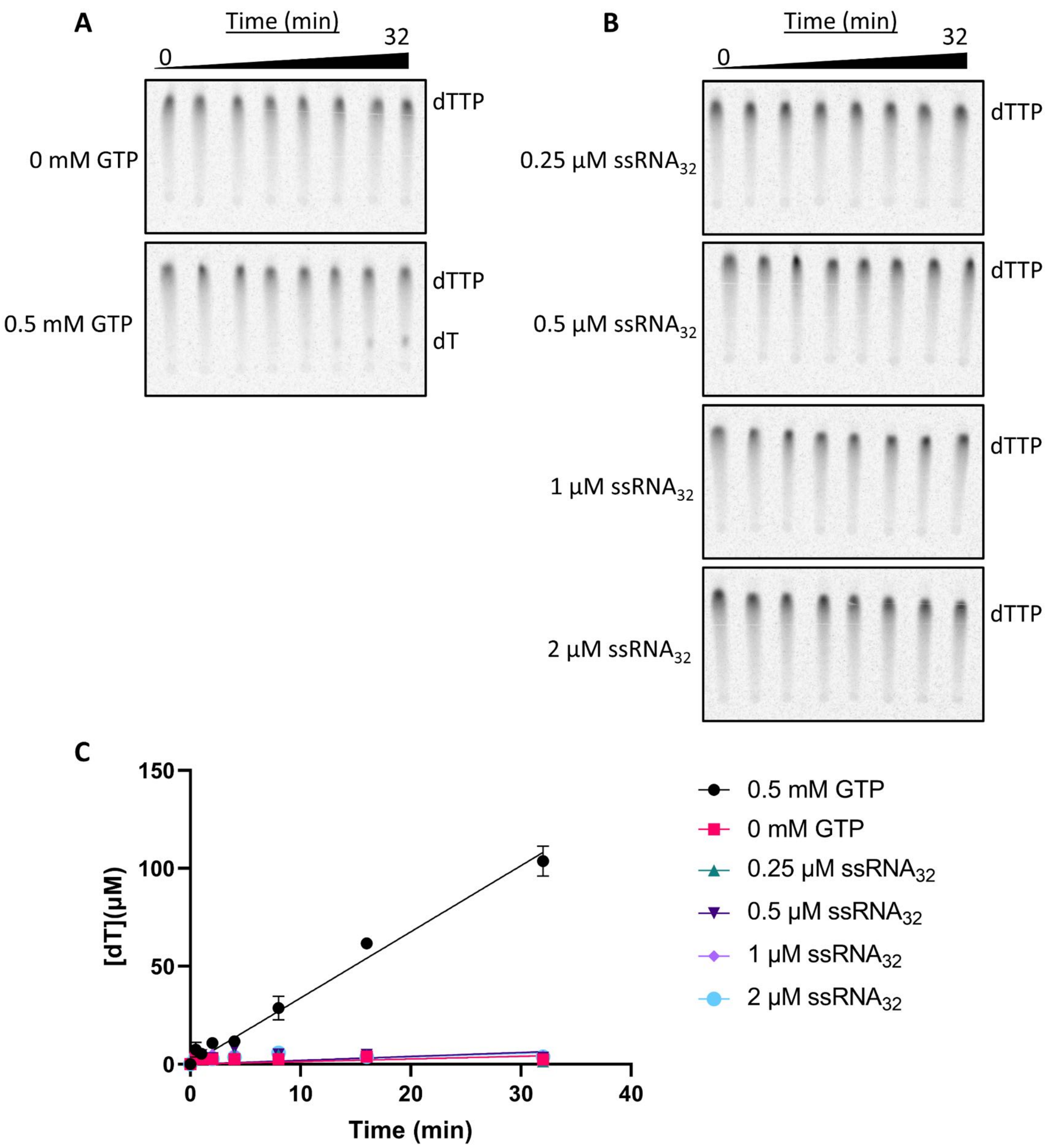

**Supplemental Figure S12. ssRNA$_{32}$ does not stimulate SAMHD1 dNTPase activity.** (**A**) Reversed-phase TLC chromatograms of the hydrolysis of dTTP (1 mM) by wild-type SAMHD1 (0.5 µM) in the absence (top) and presence (bottom, 0.5 mM) of GTP over the course of 32 minutes. (**B**) Reversed-phase TLC chromatograms of the hydrolysis of dTTP (1 mM) by wild-type SAMHD1 (0.5 µM) in the presence of various concentrations of 5'FAM-ssRNA$_{32}$. (**C**) Linear regression analysis of [dT] versus time for the reactions shown in (A) and (B). Error bars represent standard errors at each time point from three independent dTTP reactions.

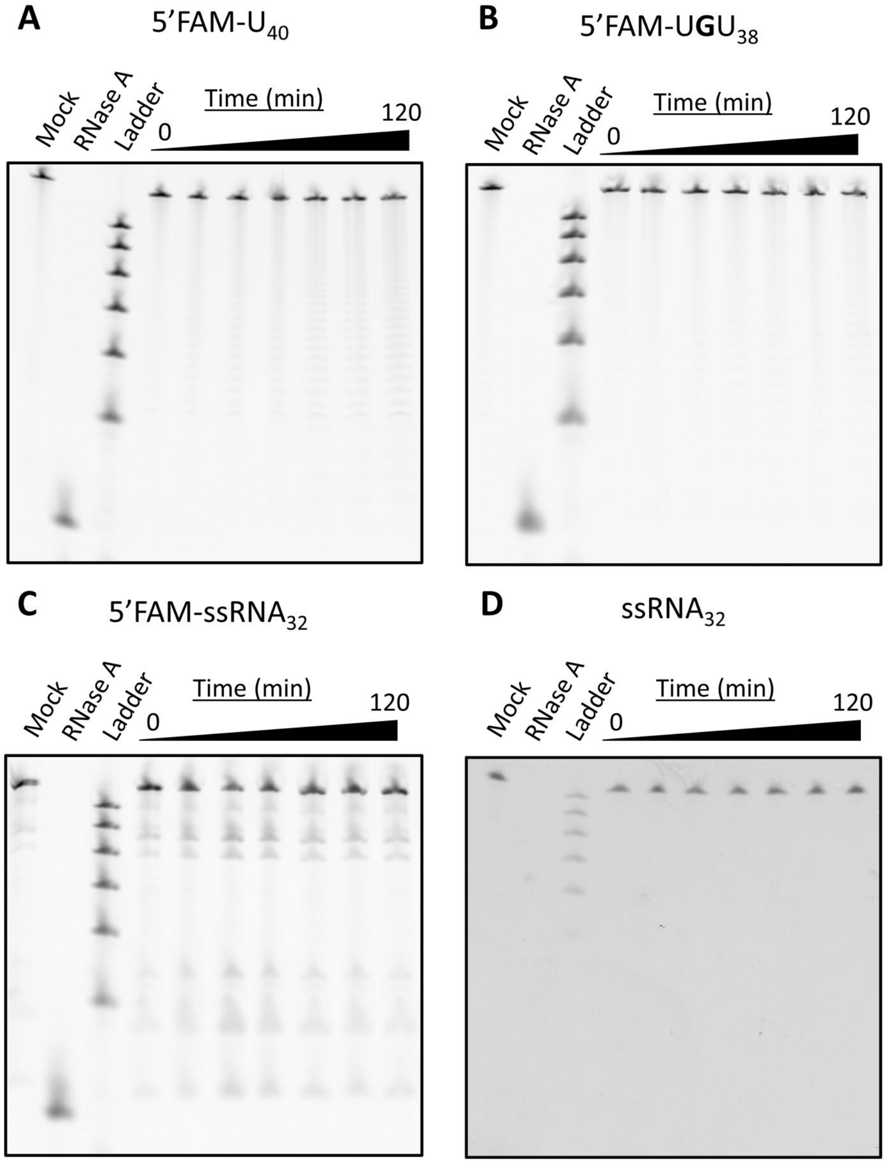

**Supplemental Figure S13. SAMHD1 possesses only trace exonuclease activity against diverse ssRNA substrates**. SAMHD1 (1 µM) was incubated with various ssRNA substrates (1 µM) for two hours. Fractions were withdrawn at regular time intervals, quenched with formamide, and ran on a 20% 19:1 acryl:bis 8 M urea gel. All reactions were run in duplicate and featured 2 hour endpoint buffer mock and RNase A positive and negative controls. (**A**) Hydrolysis reaction using 5′FAM-$U_{40}$. The gel was imaged using the fluorescence of the FAM fluorophore. The size standards are an equimolar mixture of the 5′ FAM-labeled ssDNA 30mer, 25mer, 20mer, 15mer, and 10mer used for the length dependence experiment in Fig. 2C (**B**) Hydrolysis reaction using 5′FAM-$UGU_{38}$. Gels were imaged using the fluorescence of the FAM fluorophore. (**C**) Hydrolysis reaction using 5′FAM-$ssRNA_{32}$. The gels were imaged using the fluorescence of the FAM fluorophore. (**D**) Hydrolysis reaction of unlabeled $ssRNA_{32}$ with visualization with SYBR Green II. This control establishes that the 5′FAM label is not giving rise to the negative exonuclease results.

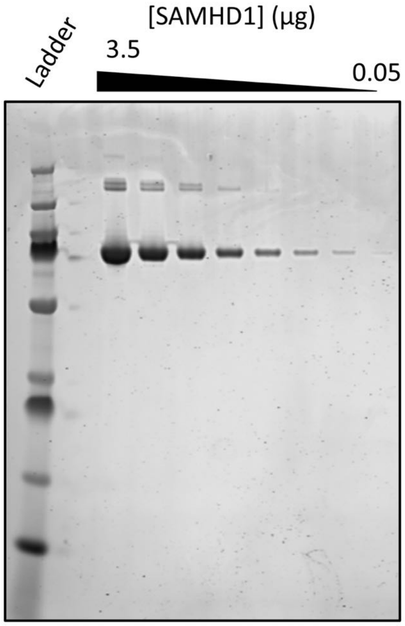

**Supplemental Figure S14. SDS-PAGE analysis of purified recombinant SAMHD1 using Coomassie fluorescence detection.** Serial two-fold dilutions of 3.5 µg purified recombinant SAMHD1 were resolved on an 4-12% gradient SDS-PAGE gel with ThermoFisher PAGEruler Plus protein ladder as a molecular weight standard. The gel was stained with colloidal Coomassie G250. The trace high molecular weight band arises from SAMHD1 disulfide linked dimers that persist after reduction with β-mercaptoethanol.

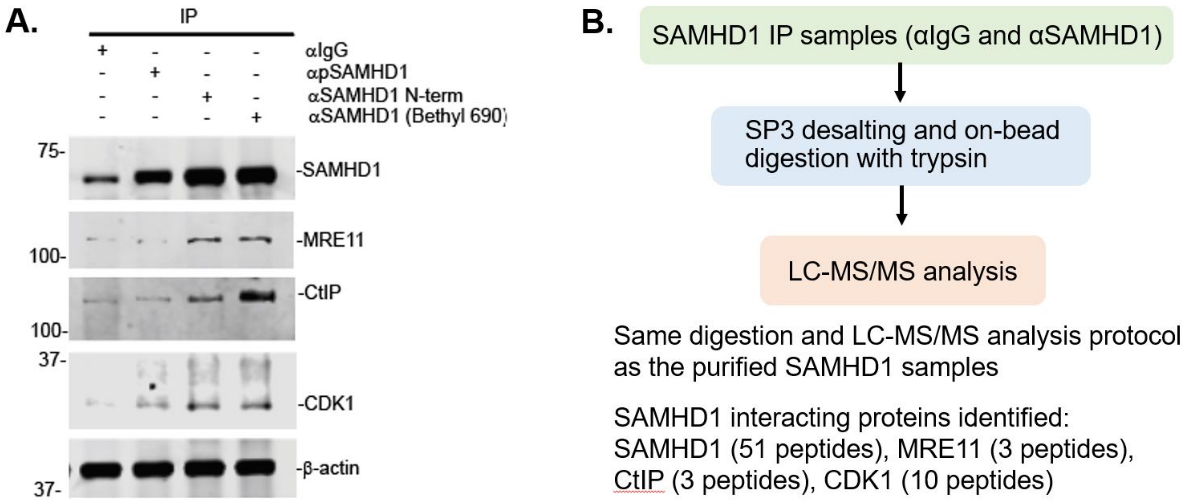

**Supplemental Figure S15. Validation of the specificity of the MS proteomics protocol using immunoprecipitation of THP1-Dual cell lysate.** (**A**) Immunoprecipitation of SAMHD1 present in THP1-Dual cell extracts co-precipitates known protein interactors MRE11, CtIP, and CDK1. (**B**) MS proteomics characterization of the SAMHD1 IP samples detects the same interacting proteins.

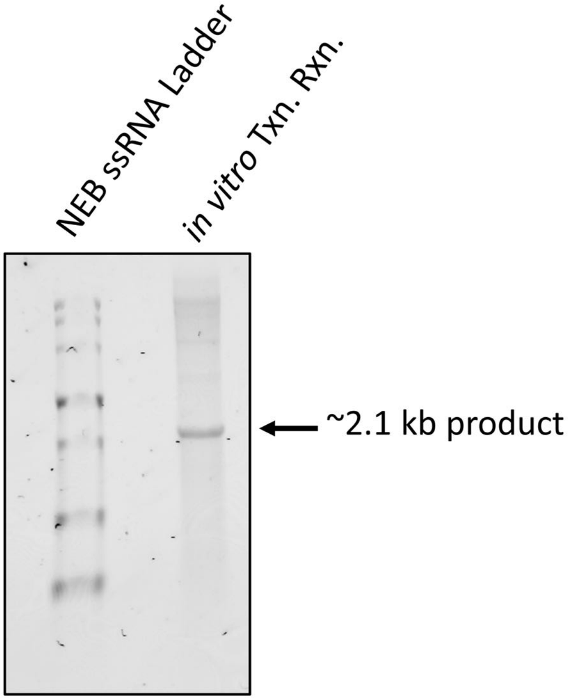

**Supplemental Figure S16. Characterization of 2kbRNA *in vitro* transcription product.** Analysis of 2kbRNA in vitro transcription product by 6% formaldehyde, 1.5% agarose gel electrophoresis. NEB ssRNA ladder was used as a standard.
